# Modeling Synaptic Effects of Anesthesia and its Cortical Cholinergic Reversal

**DOI:** 10.1101/2021.12.13.472343

**Authors:** Bolaji P. Eniwaye, Victoria Booth, Anthony G. Hudetz, Michal Zochowski

**Author notes:** Corresponding Authors: Victoria Booth, Anthony G. Hudetz, Michal Zochowski.

## Abstract

General anesthetics work through a variety of molecular mechanisms while resulting in the common end point of sedation and loss of consciousness. Generally, the administration of common inhalation anesthetics induces decreases in synaptic excitation while promoting synaptic inhibition. Animal studies have shown that, during anesthesia, exogenously induced increases in acetylcholine-mediated effects in the brain can elicit wakeful-like behavior despite the continued presence of the anesthetic. Less investigated, however, is the question of whether the brain’s electrophysiological activity is also restored to pre-anesthetic levels and quality by such interventions. Here we apply a computational model of a network composed of excitatory and inhibitory neurons to simulate the network effects of changes in synaptic inhibition and excitation due to anesthesia and its reversal by muscarinic receptor-mediated cholinergic effects. We use a differential evolution algorithm to fit model parameters to match measures of spiking activity, neuronal connectivity, and network dynamics recorded in the visual cortex of rodents during anesthesia with desflurane *in vivo*. We find that facilitating muscarinic receptor effects of acetylcholine on top of anesthetic-induced synaptic changes predicts reversal of the neurons’ spiking activity, functional connectivity, as well as pairwise and population interactions. Thus, our model results predict a possible neuronal mechanism for the induced reversal of the effects of anesthesia on post synaptic potentials, consistent with experimental behavioral observations.

**Author Summary:** Here, we apply a computational model of a network composed of excitatory and inhibitory neurons to simulate the network effects of changes in synaptic inhibition and excitation due to anesthesia and we investigate the possibility of its reversal by muscarinic receptor-mediated cholinergic effects. Specifically, we use a differential evolution algorithm to fit model parameters to match dynamics recorded in the visual cortex of rodents during anesthesia with desflurane *in vivo*. We find that changes of the fitted synaptic parameters in response to the increasing desflurane concentration matched those established by neurophysiology. Further, our results demonstrate that the cellular effects induced by anesthesia can be mitigated by the changes in cellular excitability due to acetylcholine.

## Introduction

Anesthesia is a pharmacological procedure that is used extensively in the medical profession. The goal of anesthesia is typically to suppress the patient’s conscious awareness, stress and pain associated with surgery. Several putative mechanisms have been proposed as to how anesthetic agents induce loss of awareness or consciousness, however the variety of effects of different anesthetic agents within the central nervous system make this an active area of study. Experimental studies implicate the brainstem, thalamus, and cortex as regions where neuronal activity is heavily modified by general anesthesia [1, 2]. However, the primary target region likely depends on the type of anesthetic [3]. At the single cell level, common inhalational anesthetics facilitate inhibitory transmission and suppress excitatory synaptic transmission [4, 5]. However, the extent of effects on specific synaptic receptors varies across different anesthetics (Fig 1).

**Fig 1.**
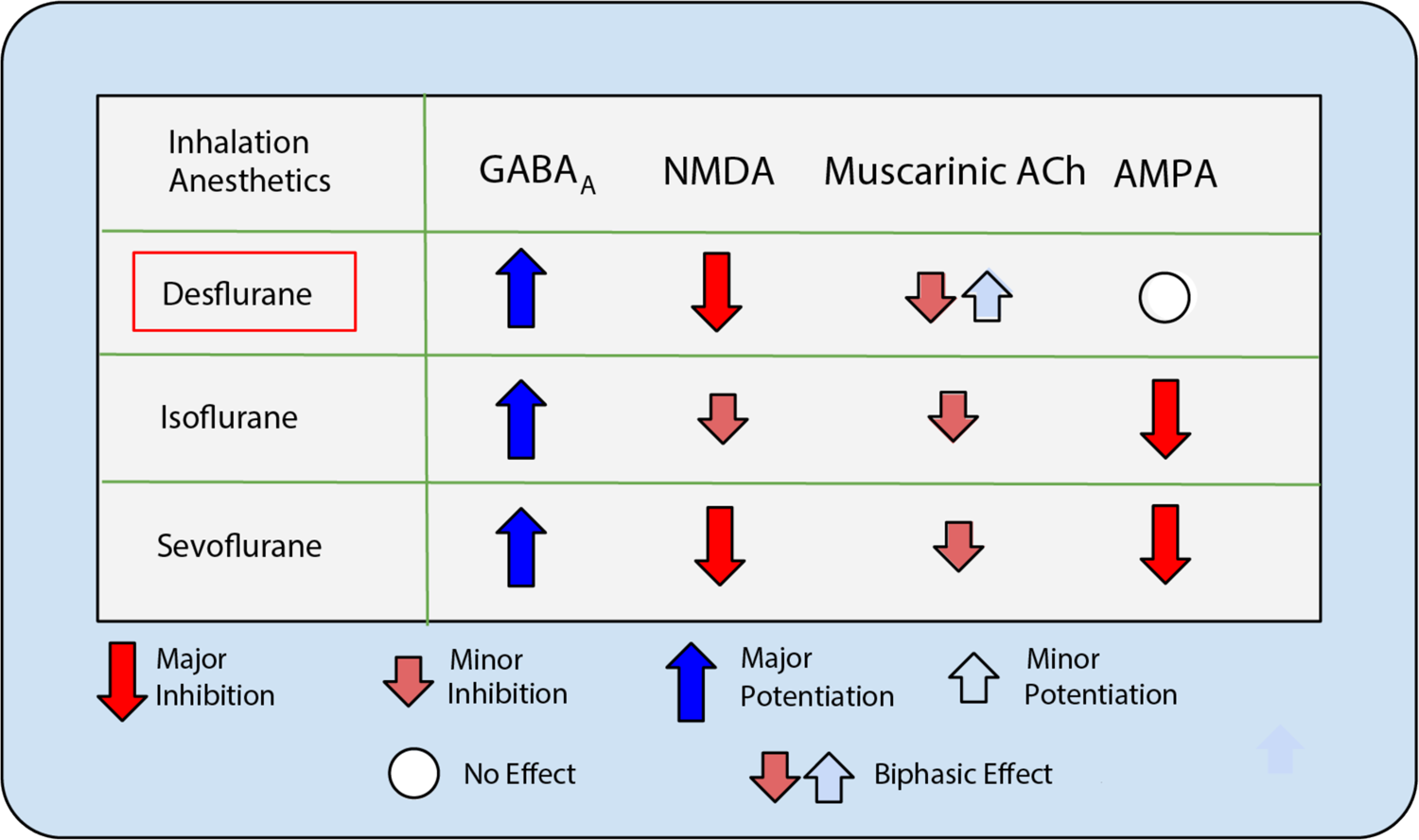
Common inhalation anesthetics have similar effects on synaptic receptors. Experimental findings show similar effects across inhalation anesthetics on synaptic receptors [19, 20]. Binding to inhibitory GABA_A_ receptors is commonly potentiated while NMDA receptor activity is commonly inhibited with the magnitude of effect varying between anesthetics. Activation of muscarinic acetylcholine receptors and AMPA receptors is inhibited by isoflurane and sevoflurane while desflurane has a biphasic effect and null effect on muscarinic acetylcholine and AMPA receptors, respectively.

Despite such differences in direct effects of different anesthetic agents, an underlying implicit hypothesis exists that there is an anesthetic agent-invariant mechanism that accounts for their final effect, the loss of awareness or consciousness. Proposed neural correlates of anesthetic action include modulation of neuronal excitability, increased network synchrony [6], disrupted brain functional connectivity and deficits in information integration [7, 8]. Integrated Information Theory is one of the leading theories of consciousness providing a general framework for how attention and awareness can be attributed to transfer and processing of information within a system [9]. Supporting this view, experimental studies have shown that information theoretic metrics of brain activity are reduced during anesthesia associated with suppressed behavioral signs of consciousness [10].

In order to understand the causal mechanisms of anesthetic action, additional experimental manipulations have been performed to modulate the state of consciousness. For example, pharmacological, electrical, and optogenetic stimulation of various brain regions have been performed to counter or reverse the unconscious state in humans and animals under the continued presence of anesthetic [11–14]. Many of these investigations utilized nicotinic [15] or muscarinic [16] cholinergic interventions. Recently, reverse dialysis delivery of the acetylcholine agonist carbachol was used to successfully reverse the effect of sevoflurane in rats *in vivo* [17]. Similar effects were observed *in vitro* when bathing cortical slices with cholinergic and noradrenergic agonists led to a reversal of slow wave oscillations induced by anesthesia [18]. These studies have shown that various behavioral expressions of the conscious state can be restored by exogenous interventions that aim to counter the pharmacological effect of anesthetics. Less investigated, however, is the question of whether the brain’s electrophysiological activity, particularly in cortical areas that are chiefly responsible for conscious representations, are also restored to pre-anesthetic levels and quality by such interventions. In other words, how do cortical neuronal activity patterns compare before anesthesia, during anesthesia and after conscious-like behavior is restored by exogenous stimulation while still in the presence of the anesthetic?

In the absence of such experimental measurements to date, computer simulations of anesthetic effects on activity in neuronal network models present a useful and promising approach. Here we embarked on such an investigation. In this study, we analyzed how single-cell synaptic effects of anesthetics translate into mesoscale changes in population dynamics that have been recorded in the visual cortex. We additionally investigated how these changes may be reversed by cholinergic activation. To do this we simulated an excitatory-inhibitory (E-I) neuron network consisting of biophysical model neurons with glutamatergic, GABAergic and cholinergic inputs to model the effects of desflurane, a common inhalation anesthetic, by varying the effect of excitatory and inhibitory neurotransmitters in a manner consistent with experimentally observed effects of desflurane at the synaptic level. To fit the model to experimentally obtained measures of *in vivo* visual cortex network firing activity at different concentrations of desflurane, we applied a differential evolution algorithm to optimize parameters modulating the effect of neurotransmitter binding at different receptors. Specifically, we quantified the graded, concentration-dependent effect of simulated anesthetic on neuronal firing rate distributions, phase coherence, monosynaptic spike transmission, network functional connectivity, and information theoretic measures of neuronal interactions, and fit these measures to corresponding experimentally measured quantities in the rodent visual cortex *in vivo*. We then used the model to simulate the presumed effect of cholinergic activation, without changing parameters for the simulated anesthetic-induced synaptic alterations, to see if these measures were reversible to near pre-anesthetic levels. Our model results provide insight into the mechanisms by which distinct neurotransmitter systems shape network behavior under the combined influence of complex pharmacological interventions that may affect the state of consciousness.

## Results

We constructed a reduced, biophysical, neuron network model to investigate how synaptic-level changes, mediated by the anesthetic desflurane, affect network-level dynamics compared to data measured in the visual cortex *in vivo*, and, separately, how cholinergic neuromodulatory changes at the cellular level may reverse these anesthetic effects. The network consisted of excitatory and inhibitory neurons interacting via synapses mediated by excitatory AMPA and NMDA receptors and inhibitory GABA_A_ receptors (see Methods section Fig 8). In addition, excitability of excitatory cells was modulated by acetylcholine (ACh), implemented via the ACh-dependent slow, hyperpolarizing K+ M-current.

We used an evolutionary algorithm (see Methods section, Fig 11) to identify optimal synaptic connectivity parameter sets (Table 1/Table 2) that most closely match multiple quantitative measures of network activity recorded under different desflurane concentrations. This allowed us to objectively find two sets of parameter modifications that fit model results to the experimental data. Namely, in one set of optimized parameters, we allowed the algorithm to optimize the inhibitory *GABA_A_* connectivity strength and excitatory NMDA connectivity strength while keeping AMPA connectivity strength constant as simulated anesthetic concentration was increased (Table 1/Table 2, A-Series). In the second set, in addition to varying the above parameters, we allowed cholinergic effects to vary with simulated anesthetic concentration (Table 1/Table 2, B-Series). The optimization was based on fitting measures of network frequency, mean phase coherence, and information theoretic measures of integration and complexity, and the parameter sets were validated using measures of synaptic connection probability and strength, as well as network functional connectivity (see Methods section, Fig 7). Optimizations were conducted separately for each anesthetic level, i.e., parameter values A1/B1 were optimized to data recorded for 0% desflurane concentration, A2/B2 for 2% desflurane, A3/B3 for 4% desflurane and A4/B4 for 6% desflurane.

**Table 1.** Parameter optimization for simulated anesthetic concentrations when performed on 10 different network realizations. A/B-series describe optimal values determined by the differential evolution algorithm fitting network connectivity parameters obtained when repeating the optimization for 10 total networks. Optimization includes A-Series, when ACh effects are assumed constant and B-Series, when ACh effects are allowed to change with anesthetic concentration. The scaling factors P_x_ scale the effects of synaptic conductances mediated by the x receptor (x = NMDA, GABA and AMPA). A1-A4/B1-B4 denote optimal parameter sets fit to experimental recordings at varying anesthetic concentrations (0%, 2%, 4%, 6% desflurane, respectively). *P_AMPA_* is only fit for the 0% anesthetic case A1/B1. Error displayed is SEM.

| | $P_{\text{NMDA}}$ | $P_{\text{GABA}}$ | $P_{\text{AMPA}}$ | $g_{\text{Ks}}$ | | $P_{\text{NMDA}}$ | $P_{\text{GABA}}$ | $P_{\text{AMPA}}$ | $g_{\text{Ks}}$ |
| --- | --- | --- | --- | --- | --- | --- | --- | --- | --- |
|  | **A-Series ( ±SEM)** |  |  |  |  | **B-Series( ±SEM)** |  |  |  |
| A1 | **1.69 ±0.05** | **3.92±0.51** | **1.27±0.67** | **0.98±0.08** | B1 | **1.69 ±0.05** | **3.92 ±0.51** | **1.27±0.67** | **0.98±0.08** |
| A2 | **1.41 ±0.03** | **4.32±0.44** | **1.27 -** | **0.98 -** | B2 | **1.63 ±0.07** | **5.51±0.33** | **1.27 -** | **1.07±0.03** |
| A3 | **1.22±0.12** | **8.13±1.21** | **1.27 -** | **0.98 -** | B3 | **1.47 ±0.06** | **6.35±0.94** | **1.27 -** | **1.23±0.02** |
| A4 | **1.09±0.09** | **10.61±1.91** | **1.27 -** | **0.98 -** | B4 | **1.36 ±0.09** | **8.17±1.41** | **1.27 -** | **1.17±0.06** |

**Table 2.** Parameter values for simulated anesthetic concentrations and cholinergic reversal results. Parameters from initial fit used to simulate anesthetic effects and cholinergic reversal. A/B-series describe optimal values of initial fit determined by the differential evolution algorithm for network connectivity parameters obtained when ACh effects are assumed constant (i.e., g_Ks_ is constant; A-Series) and when ACh effects are allowed to change with anesthetic concentration (B-Series). P_x_ denotes scaled changes in synaptic conductance’s mediated by the x receptor (x = NMDA, GABA and AMPA) as described in Table 1. A/B-Series Reversal (AR/BR series) represent simulated anesthetic reversal, obtained by increasing ACh effects (decreasing *g_Ks_* from A4/B4 levels) while keeping all other parameters constant.

|  | P<sub>NMDA</sub> | P<sub>GABA</sub> | P<sub>AMPA</sub> | g<sub>Ks</sub> |  | P<sub>NMDA</sub> | P<sub>GABA</sub> | P<sub>AMPA</sub> | g<sub>Ks</sub> |
| --- | --- | --- | --- | --- | --- | --- | --- | --- | --- |
| A-Series |  |  |  |  | B-Series |  |  |  |  |
| A1 | 1.64 | 4.51 | 1.19 | 0.94 | B1 | 1.64 | 4.51 | 1.19 | 0.94 |
| A2 | 1.46 | 5.63 | 1.19 | 0.94 | B2 | 1.58 | 5.32 | 1.19 | 1.12 |
| A3 | 1.39 | 6.72 | 1.19 | 0.94 | B3 | 1.44 | 5.81 | 1.19 | 1.24 |
| A4 | 1.26 | 7.63 | 1.19 | 0.94 | B4 | 1.34 | 7.51 | 1.19 | 1.21 |
| A-Series Reversal |  |  |  |  | B-Series Reversal |  |  |  |  |
| AR1 | 1.26 | 7.63 | 1.19 | 0.81 | BR1 | 1.34 | 7.51 | 1.19 | 1.01 |
| AR2 | 1.26 | 7.63 | 1.19 | 0.67 | BR2 | 1.34 | 7.51 | 1.19 | 0.81 |
| AR3 | 1.26 | 7.63 | 1.19 | 0.53 | BR3 | 1.34 | 7.51 | 1.19 | 0.60 |
| AR4 | 1.26 | 7.63 | 1.19 | 0.40 | BR4 | 1.34 | 7.51 | 1.19 | 0.40 |

In each optimization run we kept the network structure fixed. Particularly, when optimizing across the A-series/B-series we maintained a single network to guarantee that the cost or loss function monotonically decreased across generations. To check for robustness, we optimized parameters for 10 independent network realizations. For each network optimization, the initial pool of parameters seeding the search was kept the same. Table 1 reports mean and standard error of obtained parameter values of the 10 optimization runs. Table 2, on the other hand, represents the single optimized parameter set that was subsequently used to identify anesthetic effects on the dynamics of the network.

With synaptic connectivity parameters fixed at their levels corresponding to 6% desflurane concentration, we then simulated the reversal of the anesthetic effects by increasing AÇh effects as mediated by the muscarinic receptor dependent M-type K+ current (specifically, decreasing its conductance *g_Ks_*; Table 2, AR/BR-Series).

The synaptic connectivity parameter values determined by the evolutionary algorithm mirrored experimentally identified effects of desflurane on excitatory and inhibitory synaptic currents (See Methods section, Fig 11). For example, in the A-series parameters, there was a decrease in the effects of NMDA receptor-mediated current while there was an increase in the effect of GABA-mediated current in response to increases in anesthesia (Table 1/Table 2). A similar trend was obtained in the B-Series with the added result that decreasing effects of acetylcholine (increasing *g_Ks_*) correlated to the effects of increased anesthesia except for the change from B3 to B4. Interestingly, the optimization predicted that, in the B-Series, to offset the decrease in neuronal excitability due to decreasing ACh level (i.e., increased *g_Ks_*) with anesthetic concentration, the increase in GABA_A_ synaptic efficacy was smaller than that obtained in the A-Series, and similarly, the NMDA synaptic efficacy was systematically higher as compared to the A-Series.

Fig 2 shows example raster plots comparing experimental spike timing data collected under the varying desflurane concentrations with model results for the optimized A- and B-Series parameter sets, as well as the simulated ACh-induced reversal of anesthetic effects. The model raster plots show similar qualitative trends for increasing simulated anesthetic concentration as the experimental data, specifically spiking patterns change from asynchronous with higher spiking frequencies at simulated 0% desflurane concentration (A1/B1) to a lower frequency, more synchronized firing pattern for simulated 6% desflurane concentration (A4/B4). Furthermore, the simulated ACh reversal (AR1/BR1 – AR4/BR4) reverses those trends.

**Fig 2.**
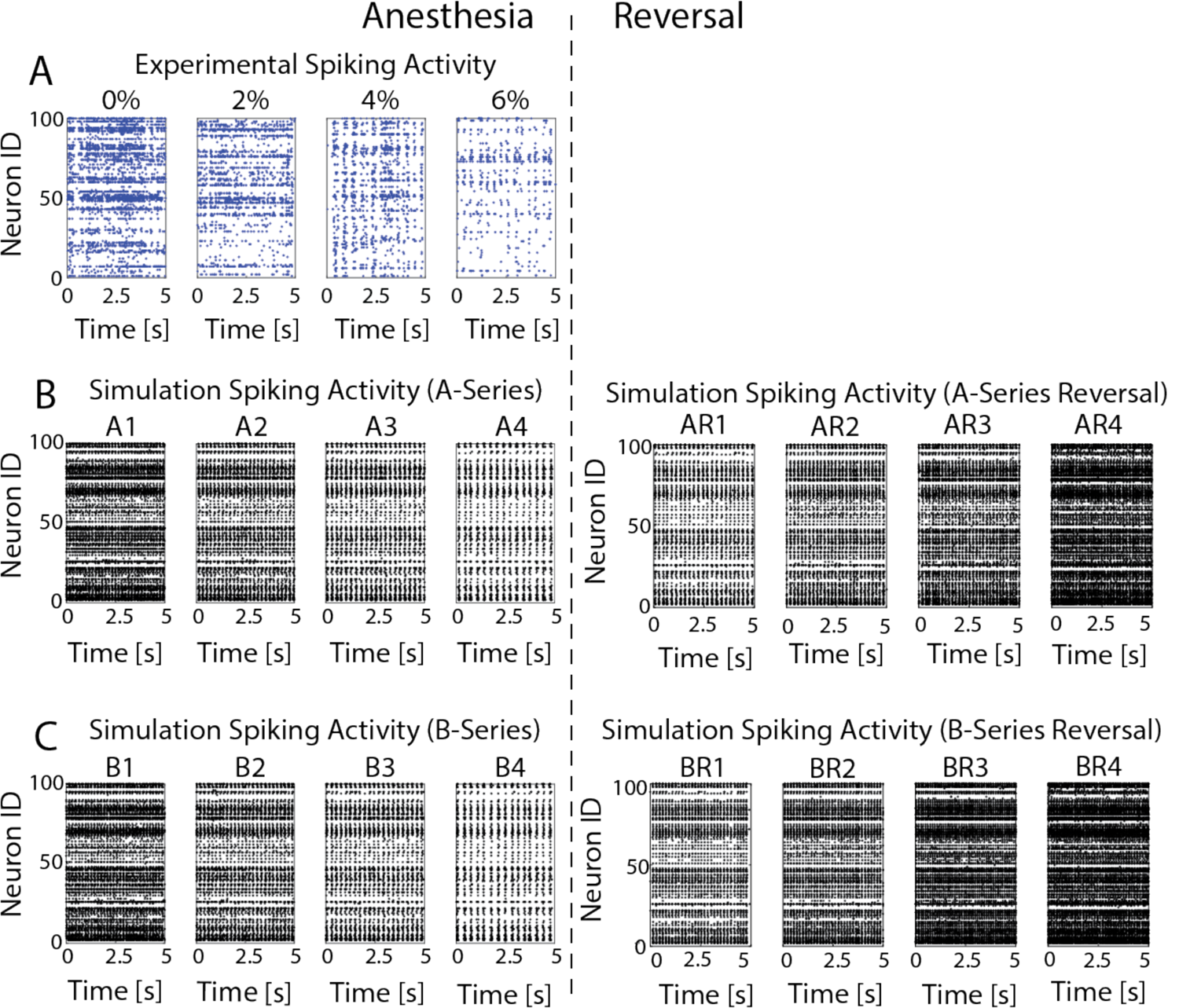
Changes in anesthesia level lead to transitions from high frequency asynchronous to low frequency synchronous spiking patterns A) Raster plots of experimentally recorded neuronal activity in response to changes in desflurane levels. For higher concentrations of desflurane (6%), oscillatory synchronous network activity can be seen in spiking dynamics. For lower levels of anesthetic, oscillations are not apparent and asynchronous activity dominates. B) Raster plots for simulated anesthetic effects in optimized model networks for constant *g_Ks_* (A series) and the simulated ACh-induced reversal of anesthetic effects (A series reversal). C) Raster plots for simulated anesthetic effects in optimized networks with changing g_Ks_ (B-series) and its reversal (B series reversal). In both B) and C), simulated anesthetic reversal shows reinstatement of asynchronous from synchronous spiking patterns.

In the following sections, we analyze how specific characteristics and measures of network dynamics, including frequency distributions and profiles, mean phase coherence, information theoretic measures, and excitatory/inhibitory connectivity probability, computed for the experimental data for progressively increased desflurane levels are reproduced in the optimized model networks. These measures were then computed for simulated increasing levels of cholinergic modulation to analyze the recovery of network dynamics during ACh-induced reversal of anesthetic effects.

### Anesthetic effects on network dynamics and their predicted ACh-induced reversal

#### Anesthetic and reversal effects on spike frequencies

We first characterized the changes in the mean neuronal spike frequency as well as the shape of the neuronal spike frequency distributions as a function of anesthetic level in the optimized model networks (Fig 3 and Fig 4A). We observed that the neurons generally fired less, in both experimental data and the simulations, as a function of anesthetic concentration. Also, the spread of neuronal firing frequencies decreased significantly with increased anesthetic level, with the loss of the right skew observed in the wake cases (0%, A1 and B1). Spike frequency decreased as a function of desflurane levels for both parameter series, (A and B series, without and with ACh changes, respectively), with a similar frequency drop, irrespective of the implemented ACh changes that affect neuronal excitability in the B series. In predicted ACh-induced reversal, the rightward skew in frequency distributions was recovered, and the B series showed stronger recovery in mean spike frequency as compared to the A series. This is because, as mentioned above, accounting for cholinergic changes on neuronal excitability under desflurane anesthesia predicts that synaptic changes are less severe. Namely, in the B series, GABA_A_ synaptic strength was not as high, and NMDA synaptic was not as low compared to the A series.

**Fig 3.**
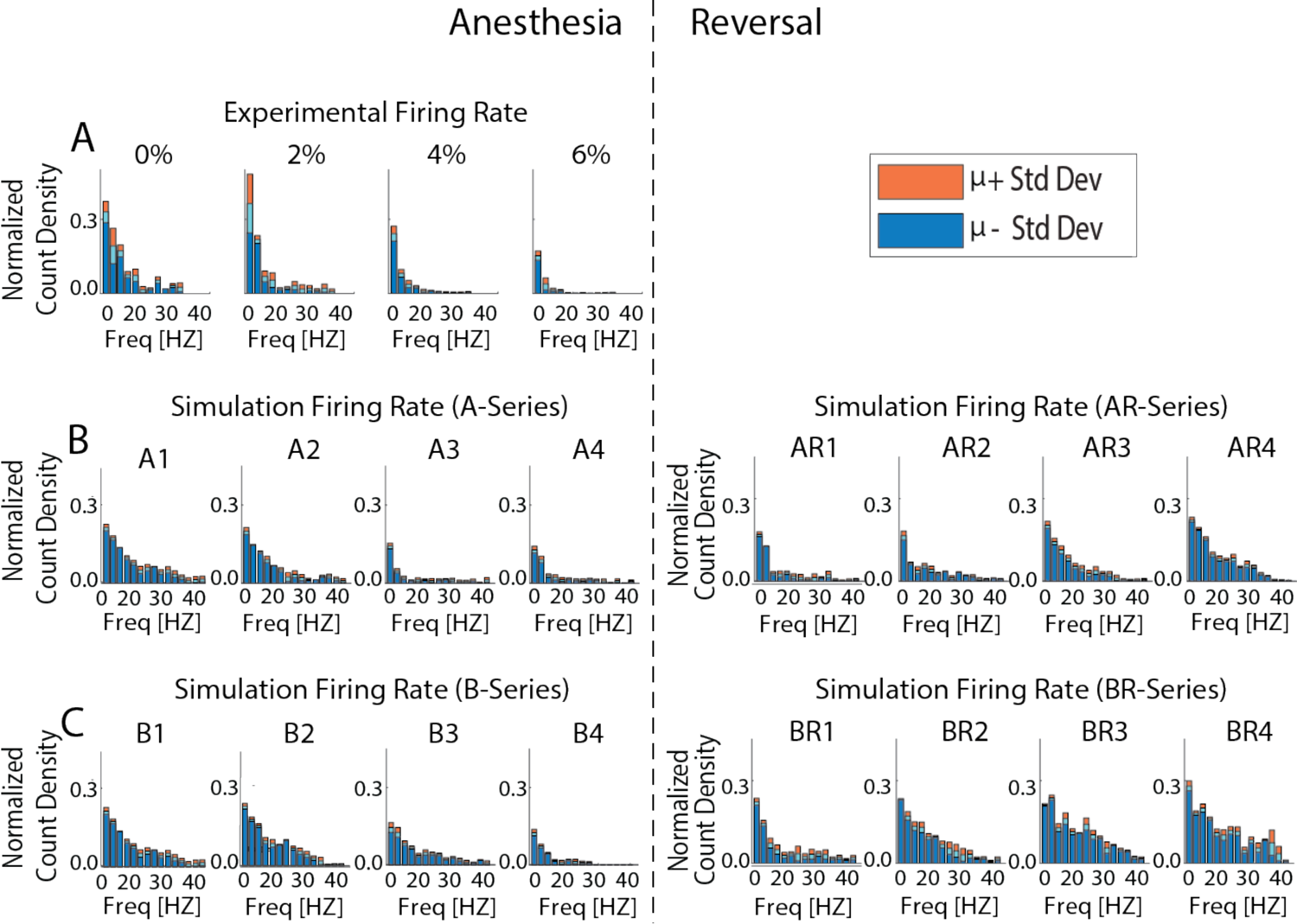
Firing rate distributions for different levels of anesthetic concentration. **A)** Changes in experimentally recorded firing rate distributions under increasing desflurane concentration (0, 2, 4, and 6%) show increased right skewness for the awake state in comparison to anesthetic states. The bins were normalized by the total number of spikes relative to the awake case (0%). **B) and C)** Firing rate distributions in optimized networks for A (B) and B (C) series parameter sets. Simulated networks show similar trends in frequency distributions when compared to experiment. The predicted ACh-induced reversal shows reinstatement of the right skew. The bins were normalized by the total number of spikes relative to the awake case A1/B1. Upper/Lower bound show histogram standard deviation.

**Fig 4.**
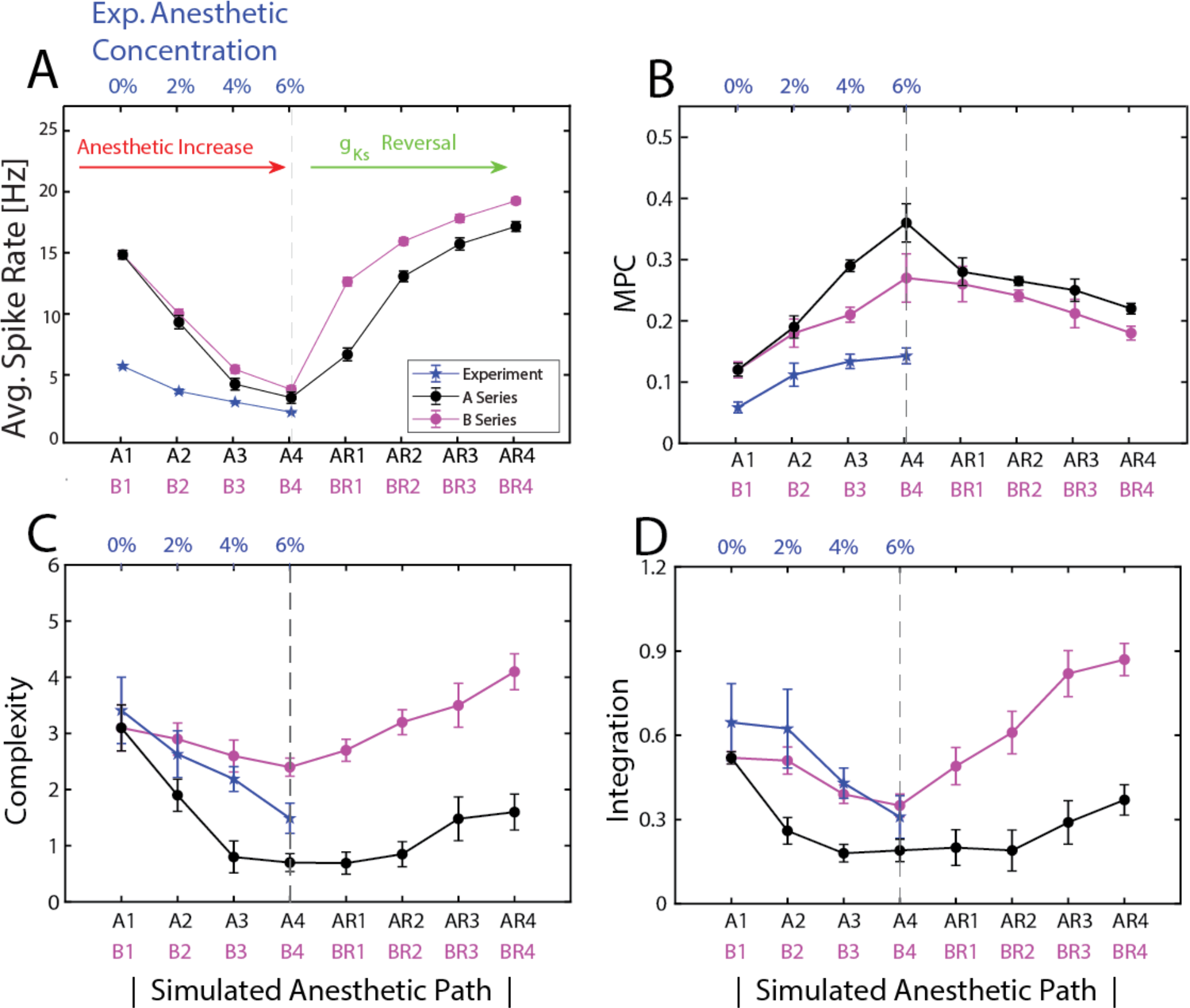
Characterization of anesthetic effects on network dynamics and their simulated ACh reversal. Measures of network dynamics computed from experimental data and optimized model networks as a function of anesthetic concentration and simulated reversal level: **A)** Average spike rate **B)**. Mean Phase coherence **C)** Complexity C(X) **D)** Integration I(X). A1-AR4/B1-BR4 (x-axis) denote simulated anesthetic concentration levels and reversal states obtained in optimized networks with corresponding parameters listed in **Table 2**. Black line denotes simulations with A-series parameter sets (g_Ks_ constant) and pink line denotes simulations with B-series parameter sets (changing g_Ks_). Blue line (with corresponding axis labels on the top) denotes measures computed from experimental spiking data at different desflurane concentrations. All calculations were made for 6s intervals and then averaged over 5 intervals. Error bars are +/-SEM.

#### Anesthetic and reversal effects on network synchrony

In both experimental and simulated results, the common feature was an increase in network synchronization as a function of increased desflurane levels. Mean phase coherence (MPC) measures the consistency of the relative phase that neurons fire with respect to each other thus taking into account non-zero time lag synchrony. For both the experimental data and the optimized network simulations, the mean phase coherence was lowest at the 0% anesthesia case and increased with increasing anesthetic concentration (Fig 4B). For higher levels of anesthesia, while the simulations and the experimental data showed increasing MPC, simulated networks exhibited a larger increase. This could be due to the fact that our model only represents local network interactions, without incorporating the existence of external inputs that could additionally desynchronize the network activity. In the visual cortex, there are non-local network inputs possibly preventing a high level of synchronization in the locally recorded network activity.

The anesthetic reversal with increased levels of ACh (i.e. decreased *g_Ks_*) led to decreases in MPC indicating desynchronization in network activity. This is not suprising, as decreased M-current leads to larger differentiation in neuronal firing rate as a function of external input and changes in the phase response curves (PRCs), which also promote desynchronisation for lower g_KS_ [21].

#### Anesthetic and reversal effects on network information metrics

We computed the information theoretic measures network integration (I(X)) and complexity (C(X)) for both experimental data and simulated network activity. Integration I(X) is a generalization of mutual information that measures the amount of total entropy of a system that is accounted for by the interactions among its elements. I(X) is zero when system elements are statistically independent [22]. Complexity C(X), on the other hand, measures the total entropy loss due to interaction of system elements, or, equivalently, the difference between the sum of the entropies of the individual elements and the entropy of the entire system. C(X) is low for systems with independent elements or with highly synchronous elements.

To compute the integration and complexity measures, 60 neurons were selected at random from both the experimental data and the simulated networks. The spike trains were then binned with 1ms bins to form binary activity vectors. I(X) and C(X) were computed by taking 3 random intervals of 6s, computing the measure on each set of intervals and then averaging the measure outcomes across the three sets (see Methods section for a detailed description). I(X) and C(X) displayed similar changes in both the experiment and simulations with increasing anesthetic concentration (Fig 4C,D). Namely, both measures decreased as a function of the anesthetic level. A difference in trends between simulation results of the A and B series is evident here, with the A series exhibiting a significantly more precipitous drop in both measures with increasing anesthetic level, as compared to the B-series. This again can be explained by the differences in network connectivity parameters (i.e., NMDA and GABA_A_ synaptic strengths) obtained for the two series. Specifically, lower NMDA synaptic efficacy and higher GABA synaptic efficacy leads to effective disconnection of the neurons in the A series networks, resulting in lower I(X) and C(X) measures.

Simulated ACh-induced reversal acted to increase both these measures (Fig 4C,D; AR/BR series). In the B series reversal, both measures recovered to values greater than the simulated waking values A1/B1. This was due to the higher NMDA and lower GABA_A_ synaptic efficacies that lead to significantly stronger excitatory interactions between the neurons in the B-series simulations, increasing I(X) and C(X) measures.

These four measures, average spike rate, MPC, I(X) and C(X), were used in the evolutionary algorithm to optimize the fit of synaptic parameters to the experimental data (see Methods section). In the next section, we validate our optimized networks by comparing additional measures of network connectivity with the experimental data.

#### Anesthetic and reversal effects on Network Connectivity

We estimated network excitatory and inhibitory synaptic strengths, as well as network excitatory and inhibitory connection probabilities, in the optimized networks and compared them directly to these same measures computed from the experimental data. These excitatory and inhibitory network connectivity measures were computed using cross correlogram analysis as described in the Methods section (Fig 10) and based on previous works [23].

The optimized networks displayed similar decreases in the strength of excitatory network connectivity with increased levels of anesthetic as observed in the experimental data (Fig 5A). Both the A-series and B-series results followed similar trajectories, with the A-series results reporting somewhat smaller excitatory connectivity strength values. This is due to the fact that the evolutionary algorithm returned significantly lower NMDA efficacy for the A-series, compared to the B-series. On the other hand, excitatory network connectivity probability is very similar for both parameter series as the structural connectivitiy density of excitatory synapses is the same in all model networks (see Methods section).

**Fig 5.**
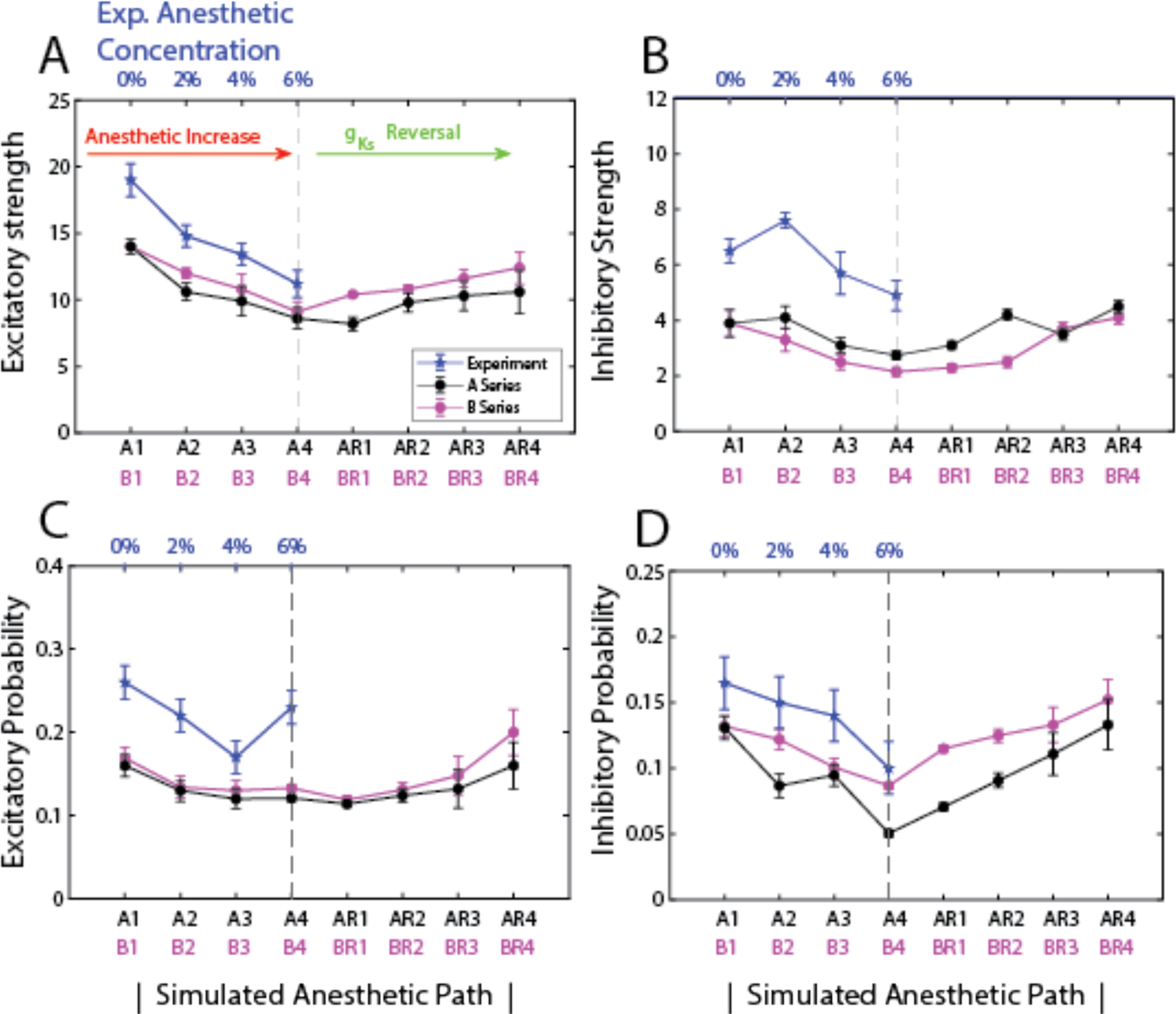
Characterization of anesthetic effects on network connectivity and their simulated ACh reversal. Measures of network connectivity computed from experimental data and optimized model networks as a function of anesthetic concentration and simulated reversal level: **A)** network excitatory connectivity strength, **B)** network inhibitory connectivity strength, **C)** network excitatory connectivity probability, **D)** network inhibitory connectivity probability. A1-AR4/B1-BR4 (x-axis) denote simulated anesthetic concentration levels and reversal states obtained in optimized networks with corresponding parameters listed in **Table 2**. Blue line (with corresponding axis labels on the top) denotes measures computed from experimental data, black (pink) line denotes measures computed from A-series (B-series) network simulations. In these measures, the presence of a significant connection was determined through cross correlogram analysis as described in Methods section. Error bars of +/-SEM.

The experimental data, as well as simulation results for both the A and B series networks, showed decreases in inhibitory network connectivity strength and probability as a function of anesthetic concentration. This seems a counterintuitive result since GABA_A_ synaptic efficacies increase with desflurane level, and were explicitly modeled as such in our networks. However, this result may be a consequence of decreases in excitatory network synaptic strength and connectivity probability. Namely, inhibitory cells receive less excitatory drive, subsequently firing fewer spikes and, thus, limiting their effect on postsynaptic targets.

Additionally, we observed that the strength of network inhibitory connectivity in the A-series networks was generally stronger than in the B-series networks. This observation agrees with the fact that the GABA_A_ conductance is higher in the A-series parameters than in the B-series. Counterintuitively, network inhibitory connectivity probability was lower and more variable in the A-series networks compared to the B-series networks.

#### Effects of ACh-induced anesthetic reversal on network functional connectivity

The results discussed above report trends observed for measures of average network activity, such as frequency, mean phase coherence, integration and complexity, as well as network connectivity strength and probability. And while ACh-induced reversal reinstated these network-level measures, the measures do not account for recovery of functional connectivity in the network which would contribute to information processing. In this section, we investigate how ACh reversal affects the relative frequency profile of individual neurons with respect to other neurons in the network and also look at effects of reversal on the cellular-level functional connectivity. These measures specifically assess whether the internal dynamic structure of network activity is reinstated during the ACh reversal.

To accomplish this, we first compared the firing rate of each neuron (or unit) in the experimental data and in the optimized networks at each level of anesthetic concentration to its firing rate in the waking state (Fig 6 and S1 Fig). In the figure panels, the x-axis represents firing frequency of individual cells for different anesthetic levels and the y-axis represents the firing frequency for the same cells in the non-anesthetic (0% or A1/B1) conditions. For the experimental data mutliple units can be potentially detected on a single electrode. This led to potential ambiguity in neurons assigned across anesthetic levels. To address the, neuron identity was based on firing rate in the 0% case. Namely, for units recorded on each electrode, the fastest firing units for 0% anesthesia were given the same ID as the fastest firing units in the 6% case. The results showing an overall linear relationship (Fig 6 and S1 Fig) indicates preservation of relative frequency ordering between the neurons. Deflection of the slope of the linear relationship towards vertical indicates the decrease in absolute firing frequency observed for different anesthetic levels.

**Fig 6.**
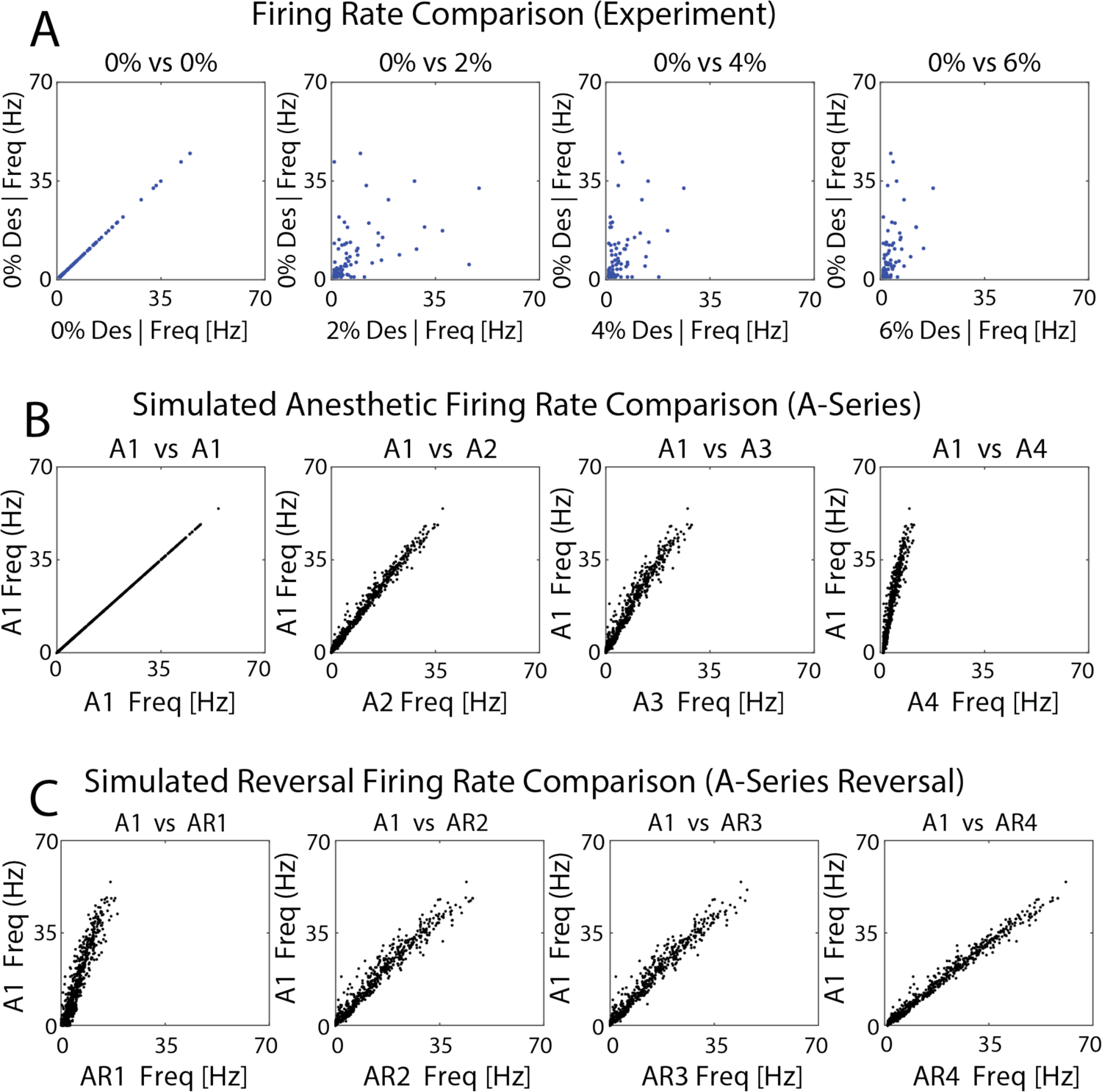
Effects of anesthetic concentration and simulated ACh-induced reversal on relative profiles of neuronal firing frequency. Each panel depicts the firing frequency of each neuron in a given anesthetic/reversal state (x-axis) compared to its firing frequency in the non-anesthetic condition (0% desflurane or A1) (y-axis) A) Units recorded in the experimental data; B,C) Neurons in A series optimized networks and reversal.

We observed that, generally, in both experiments and simulation results the relative frequency of the neurons was preserved, i.e. neurons that fired at higher frequencies as compared to other cells in non-anesthetic conditions retained higher firing frequencies at the different anesthetic levels, albeit absolute frequencies decreased. Conversely, neurons that maintained lower firing frequencies (relative to other cells) in the non-anesthetic state continued firing at lower relative frequencies in the anesthetic conditions. Qualitatively similar results were observed for A-series networks (Fig 6) and B-series networks (S1 Fig).

Importantly, during the simulated ACh-induced reversal (AR-series in Fig 6C; BR-series in S1 Fig), the relative relationship between firing frequencies of neurons remained the same, with individual cell frequencies increasing back towards their non-anesthetic values as evidenced by the slope of the linear relationship for higher reversal states tending towards one. This result suggests that individual cells return to roughly the same firing rates during ACh-induced reversal as they exhibited in the simulated waking state.

To explore detailed changes in cellular-level functional connectivity in the optimized networks, we created functional adjacency matrices from the estimated pairwise excitatory connectivity strengths at all simulated anesthetic and reversal conditions, measured via identification of the peak/trough of the spiking cross correlogram as described in the Methods section. We then calculated the cosine similarities between the created functional adjacency matrices obtained for each anesthetic and reversal level (Fig 7). A cosine similarity of 1 indicates that the functional adjacency matrices are identical, whereas cosine similarity of zero indicates that they are uncorrelated.

**Fig 7.**
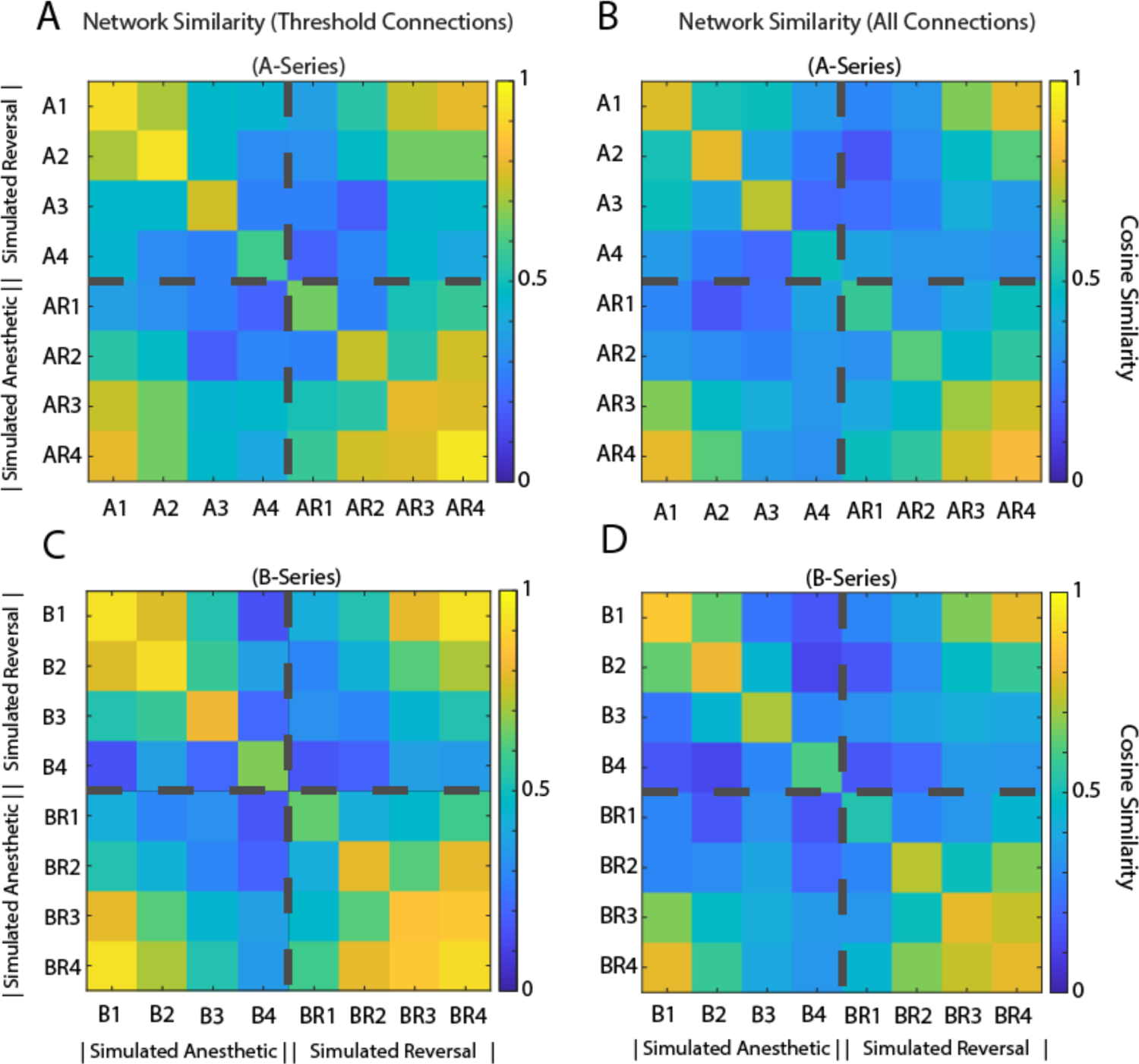
Effects of anesthetic concentration and ACh-induced reversal on the similarity between cellular functional connectivity. Cosine similarity was computed for every pairwise combination of cellular functional connectivity matrices obtained for different simulated anesthetic level (A, B - series) and reversal state (AR, BR - series). A-D) Cosine similarities of functional connectivity matrices consisting of (A, C) only significant (as detailed in the Methods section) connections and (B, D) all connections in A- and B-series network simulations. Increased similarity between AR4 and A1 (and BR4 and B1) shows reinstatement of functional connectivity that was degraded with simulated anesthetic concentrations.

The analysis was performed on all measured excitatory connections (Fig 7B, D) independently of whether their strengths passed the significance test, and separately, considering only connections that were deemed significant (thresholded, all other connections were set to zero, Fig 7A, C).

We observed that the functional adjacency matrices became less correlated with each other with increasing anesthetic levels. However, ACh reversal resulted in a significant increase in the correlation between the baseline non-anesthetic adjacency matrix (A1 or B1) and the fully reversed functional adjacency matrix (AR4 and BR4). This trend was observed when all connections were considered as well as when only significant (thresholded) connections were included, indicating that the cellular-level dynamic activity structure was largely recovered with ACh-induced reversal.

In summary, our model results showed that multiple measures of network connectivity (Figs 5 and 7) increased with ACh-induced simulated reversal suggesting that increases in cellular excitability, mediated by muscarinic effects of ACh, can reinstate network dynamics dictated by synaptic connectivity.

## Discussion

The goal of this investigation was to simulate the multisynaptic effects of an anesthetic and the modulating effect of muscarinic ACh receptor activation in a neuronal network model. We first examined if excitatory and inhibitory synaptic changes typically produced by the inhalational anesthetic desflurane led to neural network behavior similar to experimentally observed neuron activity as characterized by various measures including population firing rate, phase coherence, monosynaptic spike transmission, and the information theoretic measures integration and complexity. Second, we investigated if an exogenously induced increase in the level of ACh acting on muscarinic receptors could reverse the effect of the anesthetic as suggested by prior behavioral experiments.

### Simulation of the anesthetic effect

We simulated the effect of anesthetic desflurane on the neuronal network by reducing the response of excitatory synapses and facilitating that of inhibitory synapses. General anesthetics commonly potentiate GABAergic synaptic receptor transmission through modification of inhibitory post synaptic potential (IPSP) amplitude and duration, as well as through inhibition of glutamatergic receptor excitatory post synaptic potential (EPSP) amplitude and duration. The relative strength of these effects depends on the class of anesthetic [4, 20]. Desflurane inhibits binding at NMDA receptors while potentiating postsynaptic inhibition at GABA_A_ receptors. Some anesthetics, but not desflurane, also suppress AMPA receptors. The effect of anesthetics on nicotinic and muscarinic receptors is more diverse. Some anesthetics also modify the activity of cholinergic neurons projecting to the cortex [17]. Regarding its electrophysiological effects, desflurane has been shown to decrease average spike rate, excitatory and inhibitory monosynaptic transmission, and population measures of neuronal interactions in the cortex [8, 24]. These changes in neuronal activity observed *in vivo* have not been directly linked to the corresponding synaptic effects observed *in vitro*.

In our study we found that potentiation of inhibitory GABAergic and inhibition of excitatory glutamatergic NMDA synaptic receptors do indeed lead to graded decreases in population activity and increases in synchronization, as quantified by firing rate and mean phase coherence, as well as measured decreases in integration and complexity. Additionally, we were able to recover changes in functional network connectivity which matched changes seen in literature [25, 26]. The simulation results were robust; although some of the measures (frequency, MPC, I(X) and C(X)) were used for optimization of model parameters via the differential evolution algorithm, the results held for a wide range of non-fitted measures within physiologically reasonable limits. Moreover, the parameter fits obtained for increasing levels of anesthetic matched in their relative magnitudes to the reported anesthetic induced changes in synaptic efficacy.

### Understanding the mechanism of anesthesia through computational modeling

The cellular mechanism of anesthetic action with respect to loss of awareness has been a subject of intense investigation. Computational models are actively used to make progress in this area of research. Because differing classes of anesthetics elicit different effects on synaptic receptor subtypes, many modeling approaches aim to determine how nuanced changes in receptor binding and synaptic activity lead to changes in neural or electroencephalographic activity. For example, in mean field models, GABAergic and glutamatergic synaptic changes are attributed to a single parameter that maps to different concentrations of general anesthesia [27]. Other modelling approaches seek to understand the mechanism of specific anesthetic agents; for example, the effects of propofol have been studied through the modeling of both GABA_A_ and GABA_B_ amplitude/duration and the effects on cortical synchrony and EEG rhythms [18, 28]. Enflurane and isoflurane are other commonly modeled anesthetics where the roles of both glutamatergic receptor binding and GABAergic effects are taken into consideration [28–30]. Anesthetic action effected through post synaptic potential (PSP) changes, from a modelling perspective, is a relativity robust explanation supported by its effectiveness across modelling paradigms. These include “mean field” models as well as networks of “integrate and fire”, “Izhikevich “and “HH” neurons, which all show reduced activity and changes to population synchrony when modeling anesthetic effects on synaptic receptors [30–32].

Our study is distinguished from former computational models of anesthetic effects by the independent consideration of the effects on NMDA_R_ and GABA_R_ through PSP changes, as well as of cholinergic influence through changes in the muscarinic M-current. We also used a more biologically realistic log-normal distribution for synaptic weights [33]. Because we had access to experimental spike data, we were able to directly fit our model to empirical data at graded levels of anesthesia and then test our hypothesis regarding cholinergic anesthesia reversal.

### Anesthetic effects on spike synchrony

A common brain signature of general anesthesia is the loss of global functional connectivity between specialized regions of the cortex while local populations show increases in neural synchrony [25,34,35]. Cellular and network mechanisms leading to neural synchrony have been studied extensively in the field of computational neuroscience [36–38]. A set of possible network wide mechanisms are the PING (pyramidal interneuron network gamma) class of mechanisms, where stable, synchronous activity patterns emerge when inhibition periodically shuts down excitation in the network [39–43]. The propensity of neural network synchrony can also depend on intrinsic cellular excitability properties, an example being changes from Type 1 to Type 2 membrane excitability. Type 1 and Type 2 neural excitability describe the well-characterized differences in spike generation dynamics that can occur generally between different types of neurons, and can occur in the same neuron under different pharmacological conditions, such as changing ACh levels. Type 2 dynamics originate from increased competition between depolarizing and hyperpolarizing currents as compared to Type 1 [44]. These differences exemplify themselves in the onset and steepness of firing frequency-input (i-f) curves and the shape of phase response curves (PRCs) which in turn determine synchronizability of the networks. Neurons exhibiting Type 1 excitability respond more rapidly with higher firing frequency changes to changing stimulus magnitude as compared to Type 2 cells, and also decreased propensity to synchronize stemming from the shape of their PRC curves [21,45,46].

Thus, as also discussed below, ACh can play a double edged role in affecting network synchrony. On one hand, decreasing levels of ACh during increased anesthesia levels can promote synchrony, as it has been shown that activation of the K+ M-current mediates the transition from Type 1 to Type 2 membrane excitability [43], while on the other hand, the increase of ACh-mediated effects during reversal can offset the decreasing synaptic efficacies with higher cellular responses (increasing steepness of i-f curve). In our modelling results on ACh mediated reversal, we show that we can evoke a transition between high frequency asynchronous population behavior to low frequency synchronous activity via both mechanisms: by potentiation of IPSP and inhibition of EPSP, and ACh-mediated modulation of cell excitability. This demonstrates that it is possible for the population synchronization observed in response to anesthesia to develop in response to changes in psp alone or to concurrently active cellular mechanisms.

### Predicting anesthesia reversal by ACh

Prior experimental studies demonstrated that the behavioral expression of the anesthetic state can be reversed by stimulating the cholinergic system of the brain by various means *in vivo* and *in vitro* in both humans and animals [15,17,18,47]. To date, no modelling study has attempted to simulate the reversal of neuronal effects of anesthesia by modulating the interaction between cholinergic and other synaptic effects. In this work we demonstrated that ACh acting via blocking the muscarinic slow potassium current can reverse the general anesthetic effect on spiking dynamics and population activity, via mechanisms described above. Specifically, we showed that decreasing the influence of the M-current under simulated anesthesia leads to an increase in firing rate and neural interaction measures, showing a population wide reversal of anesthesia-induced synaptic changes. This finding suggests a possible cellular mechanism for the induced reversal of anesthesia effects on PSPs consistent with experimental studies [11, 16].

To simulate the overall effect of elevated ACh in the network, we chose to alter the muscarinic receptor-mediated pathway. The role of muscarinic ACh receptors in affecting the state of the animal depends largely on the type of general anesthetic used. Desflurane exerts a nonlinear effect on muscarinic ACh receptor activation in a concentration-dependent manner [7]. We also showed that the addition of decreasing acetylcholine influence via the muscarinic pathway during anesthesia (B series) leads to similar reversal endpoints to those with altering NMDA and GABA synaptic changes alone (A series). The choice to model changes in anesthetic ACh influence (B series) in addition to synaptic changes alone (A series) was made to generalize the effects of common inhalational anesthestics which can affect both the cholinergic as well as the glutamatergic and GABAergic pathways (Fig 1). By considering solely the effect of changes on IPSPs via GABA_R_ and EPSPs through NMDA_R_ we show that not only can changes in population activity (firing rate, synchronization and entropy), be accomplished without changes in cholinergic influence but that increasing cholinergic influence alone can reverse these effects. This demonstrates that cortical cholinergic presence has the potential to mitigate the general effects of inhalational anesthesia. In many cases, however, such as for the effects of desflurane, inhalational anesthesia can affect muscarinic and nicotinic ACh receptor binding and for this reason we decided to model the cooperative effects from changes in synaptic EPSP/IPSP and cellular excitability changes via the M-current. In the case of cholinergic reversal, however, this confounded the role of ACh, as the changes in ACh due to anesthesia could be argued to be trivially reversed in the reversal states.

In this study, we used measures of synaptic functional connectivity, computed from average pairwise correlations of neuron spiking, to quantify changes in overall network behavior in both anesthesia and reversal conditions. We showed that the cosine similarity in the functional connectivity matrix increased for the full reversal state when compared to the high anesthetic state. This means that specific neuron to neuron functional connectivity was highly correlated between the awake and reversal states but not the anesthesia states. This suggests that the functional topology of a network can be reversed through a different receptor pathway than is used to achieve the state of anesthesia. Likewise, the population measures of integration and complexity were increased by the cholinergic decrease in M-current. In fact, prior experimental studies showed that muscarinic receptor activation could reverse isoflurane-induced changes in electroencephalogram cross entropy [16]– a quantity related to brain functional complexity presumed to be associated with the conscious state [16, 48].

In the past, anesthesia reversal has been achieved by a variety of drugs and methods of administration in experimental studies. For example, microinjection of nicotine into the thalamus led to the recovery of the righting reflex in rodents anesthetized by sevoflurane [15], and a similar reversal from isoflurane was observed in response to microinjection of histamine into the basal forebrain [49]. Unlike general anesthesia, however, the mechanisms for induced reversal may be specific to the type of anesthetic agent used. An example of this can be seen when comparing the effects of the GABA_A_ antagonist, gabazine, on the effects of propofol as well as ketamine [50]. The application of gabazine led to wake-like responses when rats were sedated with propofol, which acts through potentiation of GABA_A_ receptors, but gabazine was ineffective when used during administration of ketamine, which has been known to act through modulation of NMDA receptors. These previous studies suggest that the phenomena of induced reversal can be demonstrated in controlled rodent studies, but a similar effect has been suggested in human studies [51]. Another example is the clinical case where a patient’s use of Ritalin, a central nervous system stimulant, required an increase of general anesthetic dose for sedation [52]. In rodents, Ritalin was found to cause emergence from sedation induced by isoflurane [53].

Our results predicting cholinergic recovery of neuronal population dynamics, inter-neuronal functional connectivity and complexity lends support to the evidence that the brain state altered by anesthesia is at least partially reversible. In clinical use, the effects of anesthesia can linger after the drug is no longer administered [54]. For this reason, there are both translational and phenomenological motivations to investigate induced recovery from anesthesia. Our study gives insight into the synaptic and network mechanisms by which central nervous system changes caused by anesthesia can be mitigated by the administration of a functional agonist.

### Limitations and directions for future work

We recognize a few limitations of this study. First, our model was based on random connectivity between E and I cells instead of on a detailed representation of a specific neural circuit. Other modeling studies included thalamocortical interactions [55, 56] or a cortical macro structure aimed at understanding how a multilayer architecture can influence the effects of anesthesia-induced changes in synaptic strength [55,57–59]. We argue that, for a first approximation, a generic random model is sufficient because little is known about the identity of brain regions that are responsible for mediating the anesthetic action, particularly with respect to the suppression of consciousness. We compared our model predictions to data obtained from visual cortex, which may not be the primary site of anesthetic action to suppress consciousness. It can be argued, however, that the experimentally observed changes in neural firing rates, coherence, connectivity, complexity, and other related measures are probably sufficiently general to describe the mechanism of synaptically induced network effects we intend to understand. Of course, to understand the full effect of anesthesia on the observed behavior of a live animal may require the modeling of additional effects in multiple brain regions including widespread cortical areas, thalamus, subcortical and brainstem arousal centers, to mention a few. Although other models of anesthetic mechanism have incorporated thalamocortical interactions [55, 58], none have simulated anesthetic reversal and still fall short of modeling corresponding behavioral effects. Additionally, general anesthetics have secondary effects on voltage-gated and ligand-gated channels, two-pore potassium channels, and other targets that were not represented in our model [4,7,20]. Inclusion of these additional effects could provide a more nuanced simulation with potentially closer fit to experimental data. Despite the missing details, the success of our simulations suggests that our model likely captured the essential mechanistic elements of anesthetic action and its reversal.

## Conclusion

In summary, we demonstrated that experimentally observed changes in neural activity and functional connectivity caused by desflurane could be computationally reproduced by modulating synaptic efficacy according to the known synaptic effects of the anesthetic. Additionally, we showed that by modulating the M-current alone, the effect of anesthesia on neural activity and functional connectivity in the network could be at least partially reversed. In the future, more comprehensive models that take into account cortical architecture, thalamocortical interactions and a broader array of cellular mechanisms will help to fully understand the complex roles of synaptic modulation in producing the observed neuronal network and behavioral effects of anesthesia.

## Methods

### Experimental Data

Experimental results were based on the analysis of data collected in previous studies; for an in depth description refer to the original study [60]. Briefly, rats were surgically implanted with a multishank, 64 contact microelectrode array in the visual cortex (V1). After a post-surgery recovery period, they were placed in a cylindrical anesthesia chamber for administration of inhalation anesthetic. Desflurane was applied in the sequence of 8, 6, 4, 2, and 0% inhaled concentrations for 45 to 50 min at each level.

Neural activity was recorded during the duration of the experiment and subsequently processed to extract multiunit spiking information. For this study, we analyzed unit spiking activity collected during the 0, 2, 4 and 6% desflurane exposure sessions.

### Neuron Modeling

Excitatory and inhibitory neurons are modeled using the Hodgkin-Huxley formalism [61] with parameters selected based on a model that emulated properties of both cortical pyramidal neurons and inhibitory interneurons [62, 63]. The neuron model contained sodium, delayed rectifier potassium, slow M-Type potassium and leak currents as described in the following equations:

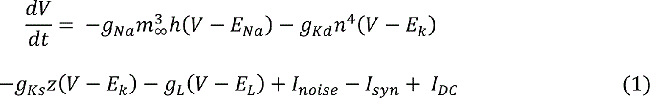

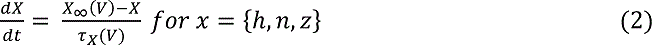

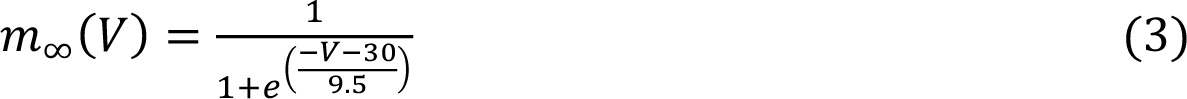

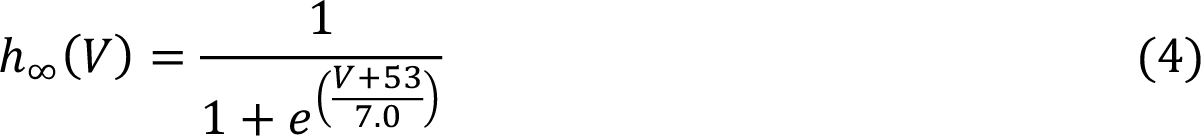

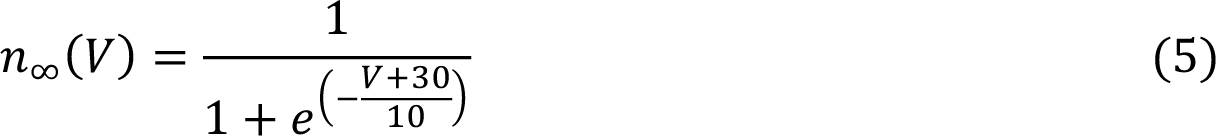

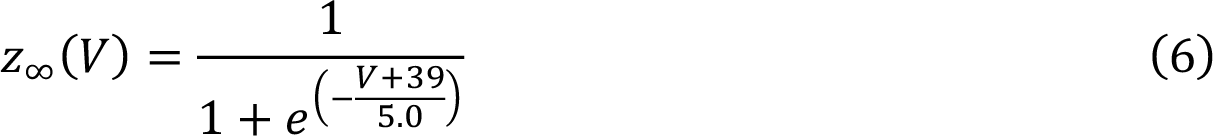

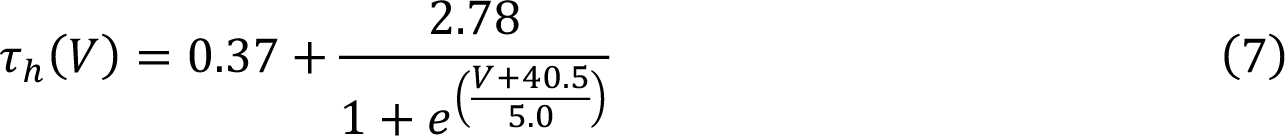

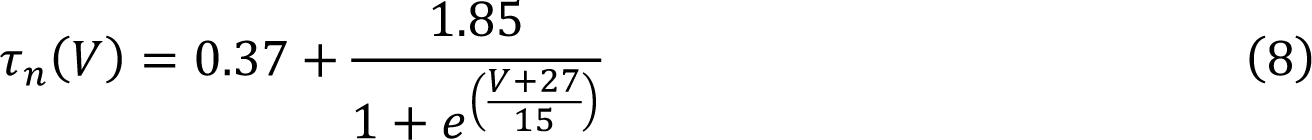

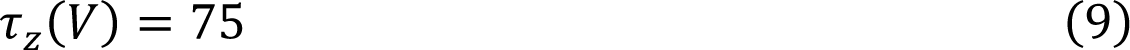

In the above, *V* is the membrane voltage while *m, n, h* and *z* represent the unitless gating variables of the ionic current conductances. *I_syn_* is the synaptic current input to the cell from other neurons in the network and has units of μA/cm^2^. *I_noise_* is a noise input consisting of randomly occurring brief current pulses with average frequency of 0.1 Hz, a duration of 2 ms and strength of 4 μA/cm^2^. This noise input was sufficiently strong to generate an action potential in the absence of any other inputs*. I_DC_* is a biasing constant current input of −0.77 μA/cm^2^. *E_Na_*, *E_K_* and *E_L_* are the reversal potentials for sodium, potassium, and leak currents, respectively, set to *E_Na_* = 55 mV, *E_K_* = −90 mV, *E_L_* = −60 mV.

This neuron model, with the slow M-type K+ current, was developed to model the muscarinic-receptor effects of acetylcholine in cortical pyramidal neurons [44]. The properties of this neuron model when *g_Ks_ = 0 mS/cm^2^* describe a neuron under high levels of acetylcholine while *g_Ks_ = 1.5 mS/cm^2^* represents a low acetylcholine state.

### Network Design

We constructed E-I networks with 800 excitatory and 200 inhibitory neurons (Fig 8B). Neurons were connected randomly with 10% probability. Synaptic strengths followed a log normal distribution, as suggested to occur in cortical networks (Fig 8A) [33]. The distribution was defined by parameters μ = −20.0, θ = 9.4, and characterized by the equation:

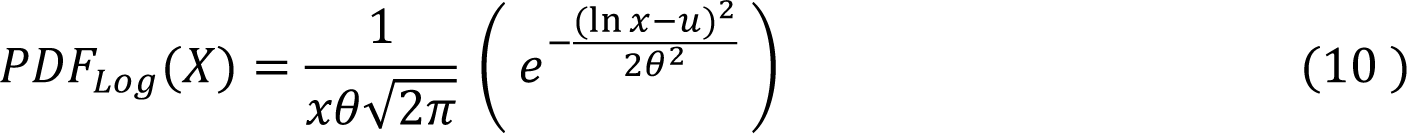

**Fig 8.**
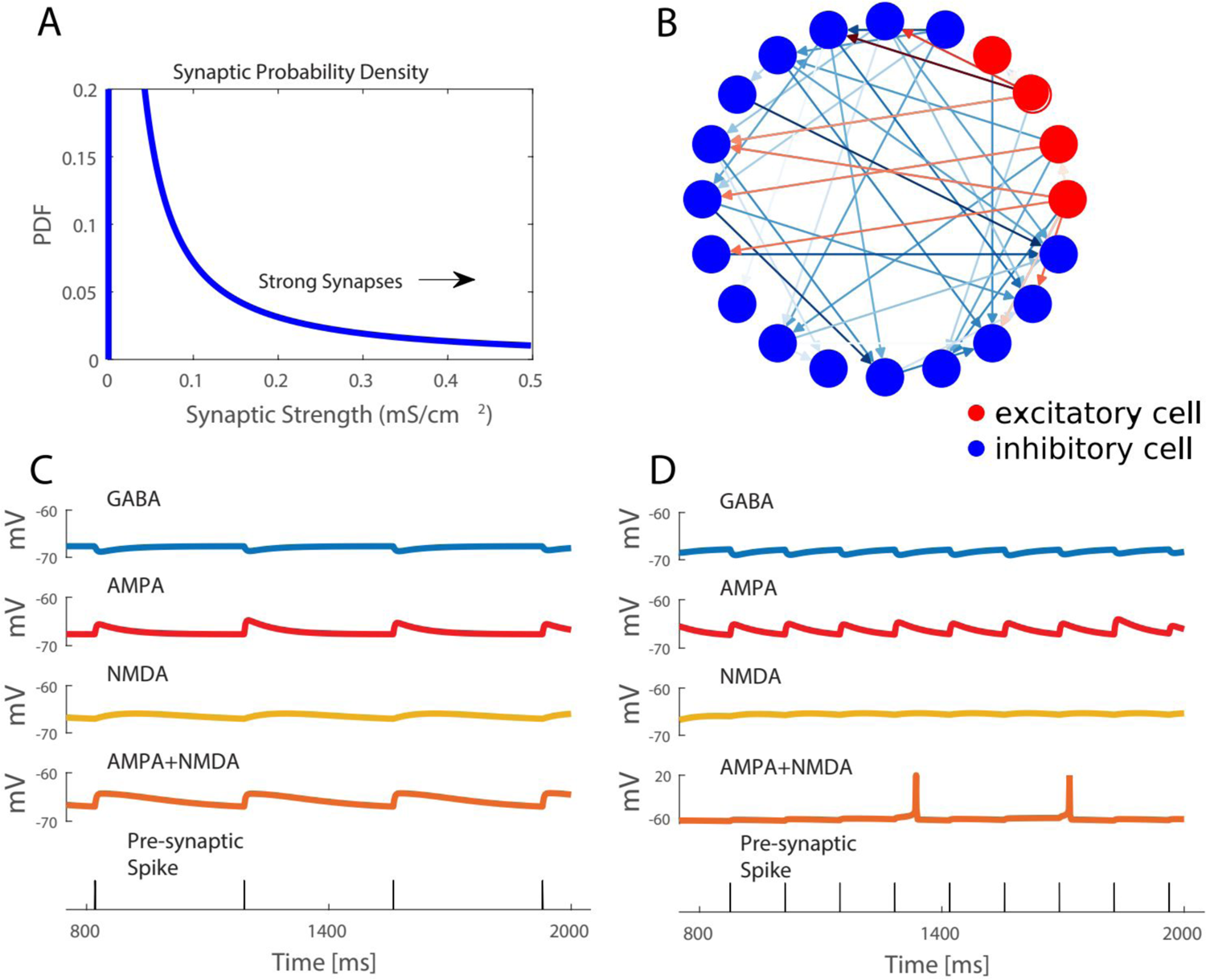
Network structure is populated by lognormal distributed random connection strengths. **A)** Synaptic strengths in model networks varied according to a lognormal distribution with a minority of connections being mediated by strong synaptic strengths, while weak synaptic strengths constitute majority of connections **B)** Simulated network consists of 200 inhibitory and 800 excitatory cells connected randomly with 10% probability. Connection color reflects the log of synaptic strength. **C, D)** Postsynaptic potential time courses in response to synaptic currents mediated by different receptors. Excitatory currents are modeled with both AMPA and NMDA mediated currents. Bottom panel shows timing of presynaptic spikes, for simplicity both inhibitory and excitatory presynaptic neurons are shown with the same spike times.

μ and θ are defined such that they are the mean and standard deviation of the logarithm of x if the logarithm of x was normally distributed. This connectivity distribution was chosen such that ∼0.2% of excitatory connections would elicit an action potential in a post-synaptic cell in the absence of other inputs for our parameter values representing the wake state. The value of 0.2% was determined by experimental data in which cross correlogram analysis showed a 0.2% “strong” connection probability among a local population of neurons [8].

Synaptic currents mediated by AMPA, NMDA and *GABA_A_* receptors were included in the network such that excitatory synaptic currents were given by *I_exc_ = I_AMPA_ + I_NMDA_* and inhibitory synaptic currents by *I_inh_ = I_GABA_*. All synaptic currents were modeled with a double exponential function of the form

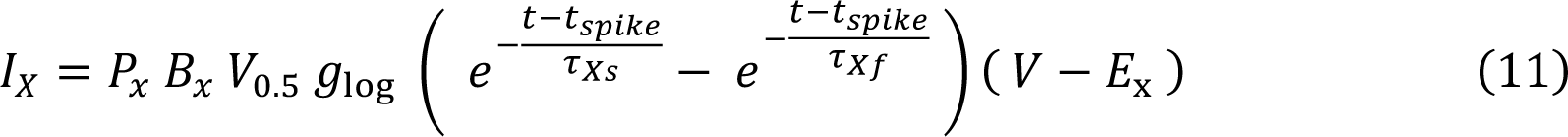

where *X* indicates the receptor type (AMPA, NMDA or *GABA_A_*), *t_spike_* is the time of the presynaptic spike and *g_log_* is the synaptic conductance drawn from the lognormal distribution. Reversal potential E_x_ was set at −75 mV for inhibitory synapses and 0 mV for excitatory synapses. The term *g_o_* will be used to refer to *B_x_ V_0.5_ g_log_*. Time constants τ_Xs_ and τ_Xf_ governed the fast rise and slow decay of the synaptic current and were set as follows:

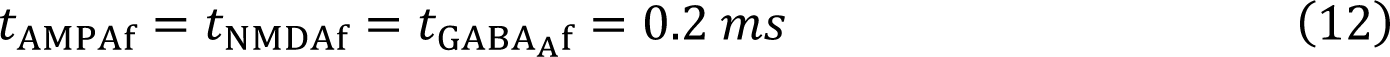

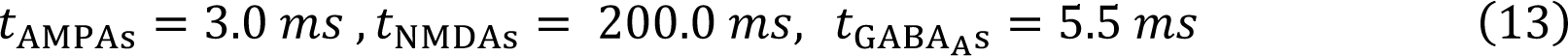

The NMDA synaptic conductance was additionally gated by the post-synaptic voltage [64, 65] described by the additional pre-factor B_x_:

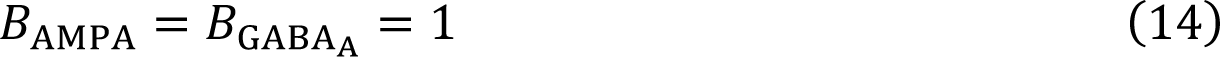

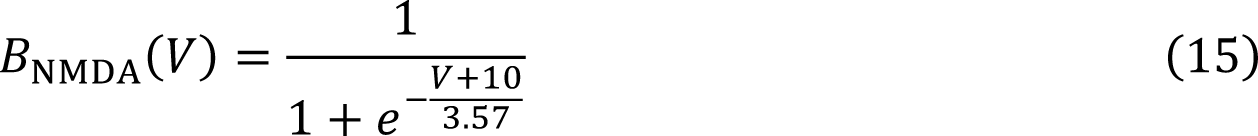

Fig 8C, D illustrates time courses of the synaptic currents. Additionally, to account for event-to-event variability, a variability pre-factor V_0.<_, randomly chosen uniformly from [0.5, 1], modulated the synaptic current induced by each pre-synaptic spike. Finally, the scaling factors *P_S_* simulated anesthetic effects on synaptic conductances. Values of *P_S_* for each receptor type were optimized to fit multiple measures of network dynamics for each level of anesthesia. Values are listed in Tables 1 and 2 that show average parameter values for optimizations performed on ten different network realizations, and the specific parameter values used for the presented analysis of results, respectively.

### Measures and Metrics

We use several different measures to quantify the changes between network states and dynamics under different levels of anesthesia observed in the experimental data and simulated in the neural network models.

#### Integration and Interaction Complexity

We computed the information theoretic measures Complexity C(X) and Integration I(X) to quantify changes in the entropy of the network [22]. I(X) is a generalization of mutual information that measures the amount of total entropy of a system that is accounted for by the interactions among its elements. I(X) is zero when system elements are statistically independent [22]. C(X) measures the total entropy loss due to interaction of system elements, or, equivalently, the difference between the sum of the entropies of the individual elements and the entropy of the entire system. C(X) is low for systems with independent elements or with highly synchronous elements.

To compute these measures, the total spiking activity from an experimental recording or a network simulation was partitioned into patterns by binning spike trains into 1 ms time bins and constructing vectors for each time bin containing a 1 at the neuron index if the neuron spiked within that time bin and a 0 if there was no spike (columns in Fig 9). The set X of unique vectors, representing patterns of spiking activity within a bin, that occurred across the data set were identified. Additionally, discretized spike vectors *X_i_, = 1, N*, were constructed for each cell (rows in Fig 9).

**Fig 9.**
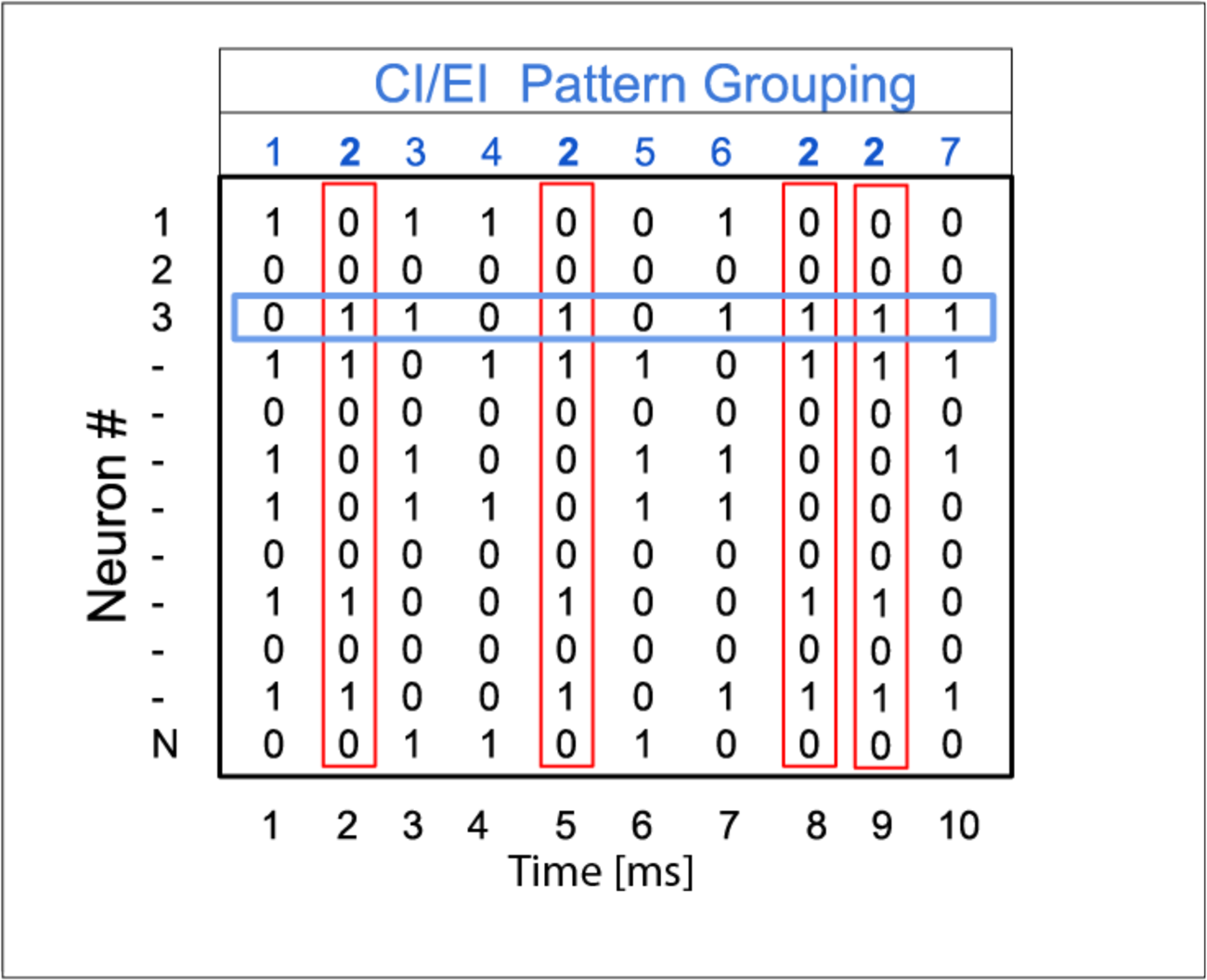
Binned spike patterns for complexity and integration measures. To compute entropy metrics complexity (C(X)) and integration (I(X)), spike trains were binned in 1 ms bins. H(X) in equation (16)/(17) is computed according to unique patterns associated with column vectors (red vectors) while H(X_i_) is the entropy associated with a single neuron spike train (blue vector).

Integration was computed as

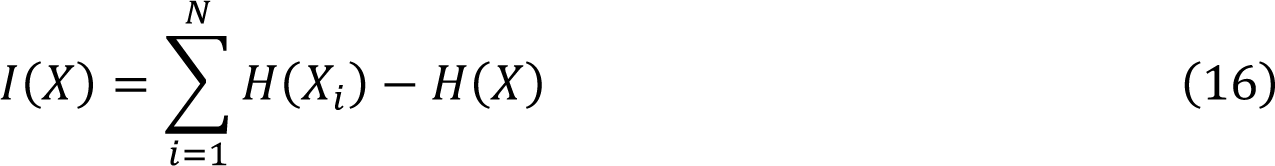

where *H(X_i_) = − ∑_k_ p_k_logp_k_* is the entropy based on the probability of a spike occurring in the *i^th^* cell, and *H(X) = − ∑_j_ p_j_logp_j_* is the entropy based on the probability of occurrence of a spike pattern vector. Complexity was computed as

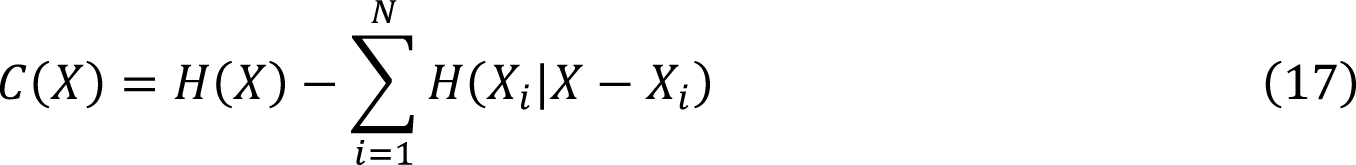

Here, *H(X_i_)* is the entropy of the spike train belonging to neuron i while H(X) is the entropy of the set of spike vector for the entire interval. *H(X_i_|X − X_i_)* is the conditional entropy where X_i_is the new spike vectors neglecting the i^th^ unit and is conditioned on the spike train of the i^th^ unit. The metric is discussed greater detail in original study [22].

#### Mean Phase Coherence

We computed mean phase coherence to quantify the average phase relation between spike times of pairs of neurons in experimental recordings and network simulation. The pairwise mean phase coherence is given by

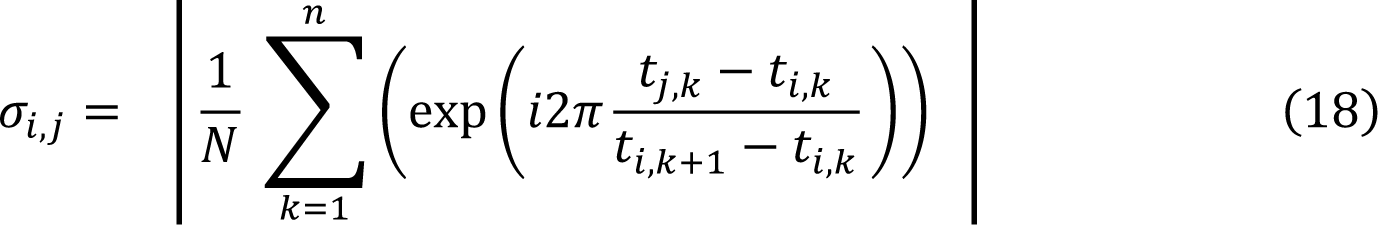

where *t(_j,k_)* is the time of the *K^th^* spike of the *j^th^* neuron and *t_i,k_, t_i+1,k_* are times of successive spikes of the i^th^ neuron. Network mean phase coherence is the average of σ_i,j_ over all pairs of neurons.

For two neurons i and j, the mean phase coherence is 1 when the spike times of neuron j always occur at the same relative phase in the cycle defined by two subsequent spikes of neuron i. Conversely, pairwise mean phase coherence is zero when spikes of neuron j occur at random phases of the neuron i spike cycle for the entire set of neurons i spike times, due to averaging of phases.

#### Functional connectivity probability and strength

Functional connectivity probability and strength were determined through cross correlogram analysis on spike trains [23] between pairs of neurons with minimum average spike rate of 1 Hz. Since experimental recordings contained on average ∼60 eligible units, these measures for the simulated networks were computed based on spike trains of 60 eligible neurons. For each pair of cells, spike trains were segmented into 40 ms intervals centered on each spike of the designated “reference” cell of the pair and discretized into 1.3 ms bins. Cross-correlations of discretized segments between the “reference” and “comparison” cell for every “reference” cell spike were summed to form cross correlograms (Fig 10).

**Fig 10.**
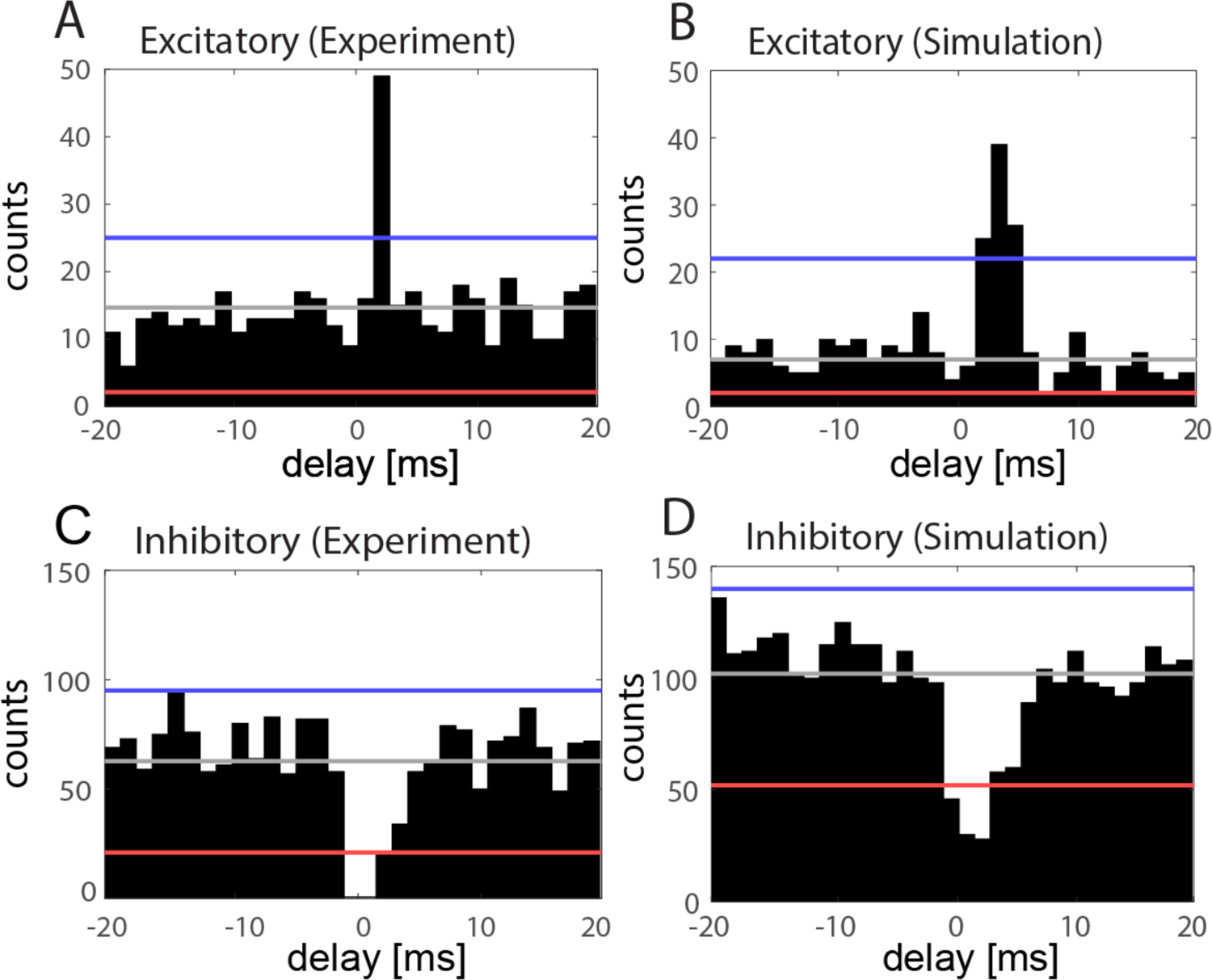
Cross Correlogram computes coincident spike relations by summing relative spike times of reference and comparison neurons A-D) Cross correlograms between example pairs of “reference” and “comparison” cells, centered at spike times of the “reference” cell, from the experimental recordings (left column) and simulated networks (right column). Significance bands were computed from a jittered data set of “comparison” cell spike times (gray line = mean of jittered data set, red line = excitatory significance, blue line = inhibitory significance, see text). A-B) Example cross correlograms showing significant excitatory connections between cell pairs. C, D) Example cross correlograms showing significant inhibitory connections between cell pairs.

Significance of correlations was determined by comparison to a constructed “jittered” dataset. The jittered data set was formed by randomly “jittering” spike times of the “comparison” cell by [−5,5] ms and then computing the cross correlogram. This was repeated by 100 times to for the jittered data set. The global confidence band for excitatory (inhibitory) connectivity was computed by taking the 97% confidence interval associated with the global peak (trough) of the jittered data set. A significant connection was determined when the peak (trough) of the original cross correlogram was greater (less) than 2 times the 97% confidence interval when measured from the mean (blue/red) [23].

Excitatory connectivity strength was determined by taking the difference in the peak height withing 0 and 5.2 ms (first four bins) and the jittered mean and dividing it by the jittered standard deviation. The inhibitory strength was computed in a similar manner by looking at the trough of the cross correlogram within 0 and 5.2 ms.

#### Parameter Optimization

Network model parameters were optimized using an evolutionary algorithm to fit measures of network frequency, mean phase coherence, integration and complexity computed from the experimental unit spiking data collected during the 0%, 2%,4% and 6% desflurane exposure sessions. The optimized parameters were the synaptic conductance scaling parameters P_AMPA_, P_NMDA_, P_GABA_ (A series) and, additionally to those, the maximal conductance of the M-type K+ current g_Ks_ (B series). The algorithm is similar to typical differential evolution procedures[66, 67]. Briefly, from a population of 30 agents (parameter sets), at each generation the 10 agents with highest cost function values were replaced with 10 new parameter sets constructed by an evolutionary algorithm described below (Fig 11A). The stopping criteria was 100 generations without change in the lowest cost function (*L(X)*) value across the population of 30 agents.

**Fig 11.**
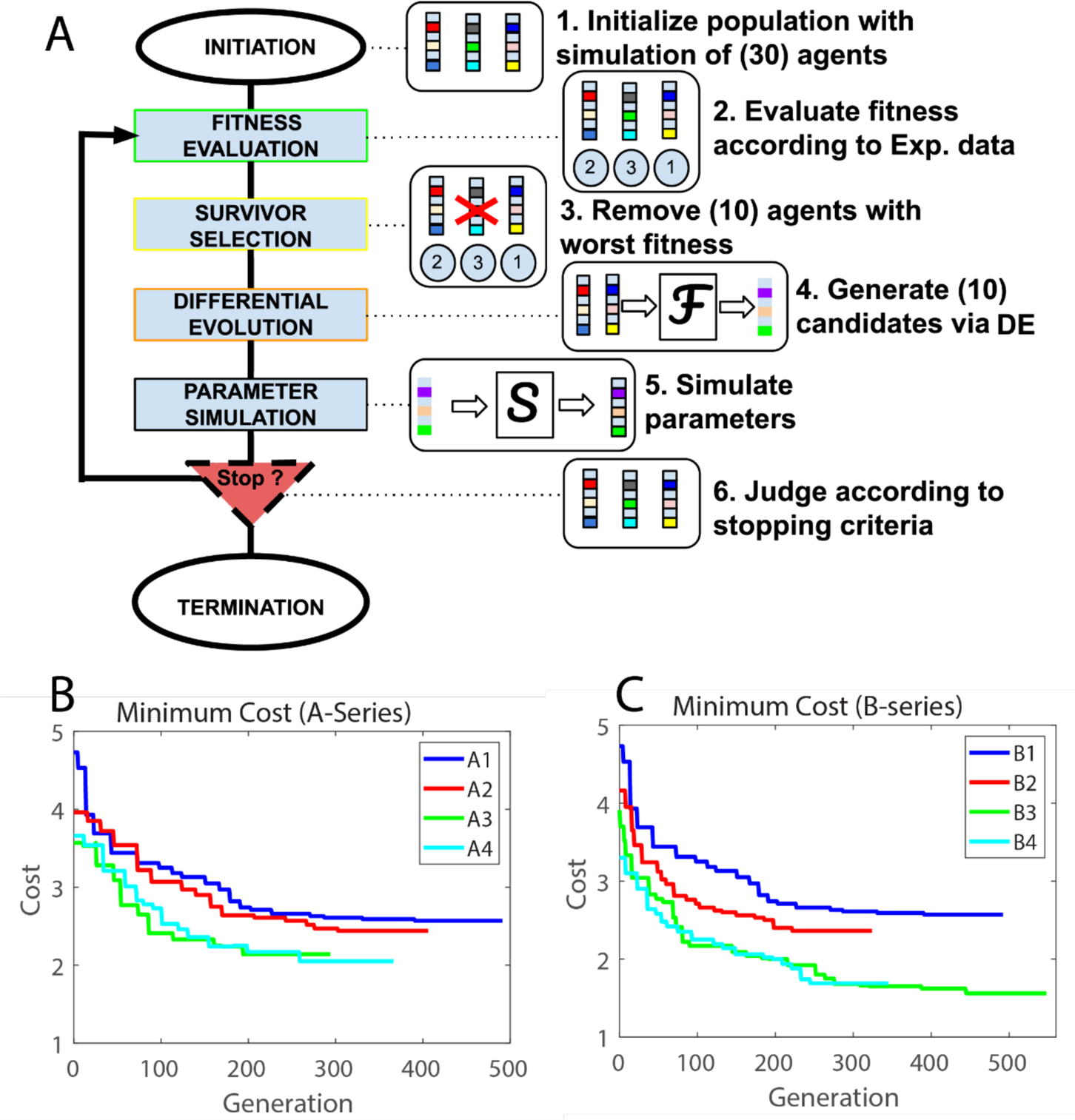
Parameter search fine-tuned through Differential Evolution algorithm. **A)** Evolutionary algorithm procedure, differential evolution, was used to optimize model parameters. For each generation, 10 agents (parameter sets) with the highest cost function from the population of 30, were chosen for replacement. Algorithm was repeated until stopping criteria of 100 generations without change in lowest cost function value across the population was met. **B,C)** Lowest cost function values across the parameter set populations at each generation for the A-series (B) and B-series (C) parameter optimizations. Population A1/B1, A2/B2, A3/B3 and A4/B4 were optimized to experimental data from the 0%, 2%, 4% and 6% anesthetic cases, respectively. The optimizations for A1 and B1 were identical. In the A-series (A2-A4), *P_NDMA,_ P_GABA_,* were optimized and in the B-Series (B2-B4), *P_NDMA_, P_GABA,_ g_Ks_* were varied.

The initial population of 30 parameter sets representing the 0% anesthetic state was chosen from the 256 parameter sets generated by assigning parameter values from the following sets: P_AMPA_, P_NMDA_ ∈ {0.5, 1.0, 1.5, 2.0}, P_GABA_ ∈ {2.5, 5.0, 7.5, 10.5} and g_Ks_ ∈ {0.3, 0.7, 1.1, 1.5} mS/cm^2^. Model networks with fixed connectivity structure and synaptic strength g_0_ values were simulated with each parameter set for 20 s and frequency, mean phase coherence, integration, and complexity measures were computed based on spiking activity excluding the initial 1s, to avoid initial transients. The cost or loss function, C(X), based on these measures, x = frequency, MPC, I(X) and C(X), compared values computed from simulations, x_sim_, and experimental data, x_exp_, at 0% anesthetic state as follows:

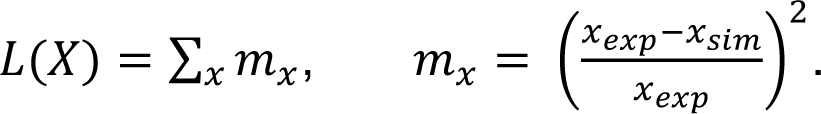

The 20 lowest cost parameter sets were kept and each parameter value was randomly varied uniformly by 10% of its value to avoid duplicate values. The final 10 parameter sets were then constructed using the differential evolution algorithm.

Similar to typical differential evolution procedures [66, 67] we set a cross over probability CR = 0.8 and a differential weight DW = 2. From the subpopulation of 20 parameter sets, 10 randomly chosen sets, *a^k^ (k = 1, …, 10)*, formed the basis for 10 newly created sets, *e^k^ (k = 1, …, 10)*. For each set *a^k^*, 3 different sets *b^k^, c^k^* and *d^k^* were chosen that were different from a^k^ and each other. Then, for each element i = 1, …, 4 in the set, a random number ρ_i_ from the uniform distribution [0, 1] was chosen. If ρ_i_ was less than CR, a new parameter value *e^k^_i_* was generated as 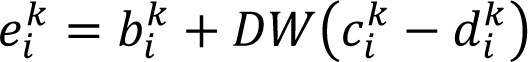; otherwise *e^k^_i_ = a^k^_i_*.

This was done for each element in the new agent and was repeated until 10 new agents were created. After this was done the 10 new agents were simulated and then the 30 total parameters were evaluated for their cost. The 10 with the highest cost (worse fit) were then rejected and the process was repeated.

We performed 2 parameter optimizations, A-Series and B-Series, to parse out potentially different effects of anesthetic modulation on synaptic conductances only (A series) and of combined modulation on synaptic conductances and cholinergic effects (B series) (Fig 11 B,C). In both scenarios, populations A1/B1 were the result of optimizing P_AMPA_, P_NMDA_, P_GABA_, g_Ks_ to the experimental 0% anesthetic case. In the A-series, P_NMDA_, P_GABA_ were optimized while in the B-series, P_NMDA_, P_GABA_, g_Ks_were optimized to the 2%, 4% and 6% anesthetic cases. Optimizations for the 6% anesthetic case, A4/B4, were initiated from parameter values constrained by experimental reports of 20% average decrease in NMDA-mediated synaptic signaling and 40% increase in GABA-ergic synaptic signaling under desflurane [68, 69]. These initial values were randomly varied uniformly by +/- 5% to generate variability in the event of parameter convergence. In the optimizations for the 2% and 4% anesthetic cases, A2/B2 and A3/B3, respectively, the initial population for A2/B2 was A1/B1, and the initial population for A3/B3 was A4/B4.

### Simulation of ACh reversal

To validate robustness of the parameter optimization, we ran our optimization for 10 network realizations, keeping the network structure fixed for all anesthetic levels. The average and error (SEM) for the optimized parameters across these 10 networks is shown in Table 1. Table 2 lists the parameter values with the lowest cost function for one of these optimization runs that we used in our model analysis. Simulated cholinergic reversal (AR1-AR4/BR1-BR4) was modeled by decreasing the value of g_Ks_ from the values in A4/B4 to 0.4 mS/cm^2^ such that there were 4 values in the reversal series.

### Simulations

Custom C++ code was developed for numerical simulations which was run on the Greatlakes High Performance Cluster. For the evolutionary algorithm each model simulation was run for 20s. The stopping criteria was met when the lowest minimum cost remained unchanged for 100 generations. To check the robustness of the current parameter set, 10 additional generations were run with model simulations of 80s and an increased crossover probability (CR=0.9). We detected no change in the minimum cost parameter set. For the results shown in Figs 3-5 each simulation was simulated for 150000 ms or 150 s. The length of this runtime was necessary to result in enough spike times to calculate metrics based on cross correlograms. Results in Figs 4 and 5 are for 10 simulation runs in which network connectivity was randomized across runs but maintained for the different simulated anesthetic levels. In this way, each of the 10 simulation runs corresponds to a unique simulated experiment. On each run the voltage and gating variables were subject to random initial conditions independent of the network seed. On initialization V was uniformly varied between [−72,−32] mV, n between [0.2,0.6], z between [0.2,0.3] and h between [0.2, 0.6] while m was initialized at 0 for all runs. The equations were integrated using the 4^th^ order Runge Kutta method.

## Acknowledgements

Research reported in this publication was supported in part by the National Institutes of Health, National Institute of General Medical Sciences R01-GM056398 (AH), and National Institute of Biomedical Imaging and Bioengineering R01EB018297 (MZ, VB), by the National Science Foundation BCS-1749430 (VB, MZ), and by the Rackham Merit Fellowship (BPE). The content is solely the responsibility of the authors and does not necessarily represent the official views of these funding agencies.

## Supporting Information

**S1 Fig. B-Series effects of anesthetic concentration and simulated ACh-induced reversal on relative profiles of neuronal firing frequency.** Each panel depicts the firing frequency of each neuron in a given anesthetic/reversal state (x-axis) compared to its firing frequency in the non-anesthetic condition (B1) (y-axis) A,B) Neurons in B series optimized networks and reversal.

